# Planktonic protist diversity across contrasting Subtropical and Subantarctic waters of the southwest Pacific

**DOI:** 10.1101/2021.09.12.459994

**Authors:** Andres Gutiérrez-Rodríguez, Adriana Lopes dos Santos, Karl Safi, Ian Probert, Fabrice Not, Denise Fernández, Priscillia Gourvil, Jaret Bilewitch, Debbie Hulston, Matt Pinkerton, Scott D Nodder

## Abstract

Planktonic protists are an essential component of marine pelagic ecosystems where they mediate important trophic and biogeochemical functions. Although these functions are largely influenced by their taxonomic affiliation, the composition and spatial variability of planktonic protist communities remain poorly characterized in vast areas of the ocean. Here, we investigated the diversity of these communities in contrasting oceanographic conditions of the southwest Pacific sector (33-58°S) using DNA metabarcoding of the 18S rRNA gene. Seawater samples collected during twelve cruises (n = 482, 0-2000 m) conducted east of New Zealand were used to characterize protist communities in Subtropical (STW) and Subantarctic (SAW) water masses and the Subtropical Front (STF) that separates them. Diversity decreased with latitude and temperature but tended to be lowest in the STF. Sample ordination resulting from the abundance of amplicon single variants (ASVs) corresponded to the different water masses. Overall, *Dinophyceae* (34% of standardized total number of reads) and Chlorophyta (27%) co-dominated the euphotic zone, but their relative abundance and composition at class and lower taxonomic levels varied consistently between water masses. Among Chlorophyta, several picoplanktonic algae species of the *Mamiellophyceae* class including *Ostreococcus lucimarinus* dominated in STW, while the *Chloropicophyceae* species *Chloroparvula pacifica* was most abundant in SAW. *Bacillariophyta* (7%), *Prymnesiophyceae* (5%), and *Pelagophyceae* (3%) classes were less abundant but showed analogous water mass specificity at class and finer taxonomic levels. Protist community composition in the STF had mixed characteristics and showed regional differences with the southern STF (50°S) having more resemblance with subantarctic communities than the STF over the Chatham Rise region (42-44°S). Below the euphotic zone, Radiolaria sequences dominated the dataset (52%) followed by *Dinophyceae* (27%) and other heterotrophic groups like Marine Stramenopiles and ciliates (3%). Among Radiolaria, several unidentified ASVs assigned to *Spumellarida* were most abundant, but showed significantly different distribution between STW and SAW highlighting the need to further investigate the taxonomy and ecology of this group. This study represents a significant step forward towards characterizing protistan communities composition in relation to major water masses and fronts in the South Pacific providing new insights about the biogeography and ecological preferences of different taxa from class to species and genotypic level.

**Highlights:**

- Water-mass preference of different taxa emerged at class, species and genotypic level.
- *Mamiellophyceae* green algae dominated in subtropical waters.
- *Dinophyceae* and *Chloropicophyceae* green algae dominated in subantarctic waters.
- A diverse assemblage of Radiolaria dominated the mesopelagic zone.
- Small rather than large taxa dominated phytoplankton blooms in subtropical waters.

## 1. INTRODUCTION

Planktonic protists, including phototrophic, heterotrophic and mixotrophic single-celled eukaryotes, have key roles in the functioning of marine ecosystems (Caron et al. 2012). Phytoplankton are responsible for 50% of global primary productivity (Field et al. 1998). Most of this primary production is consumed and processed by heterotrophic protists (i.e. microzooplankton) before becoming available for larger zooplankton and higher trophic levels (Calbet and Saiz 2005; Calbet and Landry 2004; Zeldis and Décima 2020). From a biogeochemical perspective, the microbial production, consumption and remineralization of organic matter is at the core of global biogeochemical cycles including the nitrogen and carbon cycles, and is pivotal in regulating the ocean’s capacity to sequester atmospheric CO_2_ via the biological carbon pump (Boyd et al. 2019; Turner 2015).

The trophic and biogeochemical processes driven by microbial communities are influenced by their taxonomic composition, which is tightly coupled to physico-chemical conditions. With increasing evidence of climate change effects on the physico-chemical status of the ocean (e.g. warming, increased stratification and reduced nutrient supply, and acidification) (Henley et al. 2020; Pörtner et al. 2014; Sarmiento et al. 2004) it becomes imperative to better characterize the biogeography and distributions of microbial communities in relation to oceanographic provinces (Cavicchioli et al. 2019). This is required to establish a conceptual framework and baseline upon which future environmental change can be evaluated.

The diversity and dynamic nature of microbial communities has precluded a comprehensive characterization of species composition and distributional patterns across at relevant temporal and spatial scales (Wietz et al. 2019). Extensive application of DNA metabarcoding approaches during the last 10 years have contributed significantly to this end by characterizing the diversity of marine protist communities over a wide range of temporal and spatial scales with unprecedented taxonomic resolution and coverage (Santoferrara et al. 2020). Despite these efforts there are still vast ocean regions like the southwest (SW) Pacific Ocean that due to its large size and remoteness remain largely unexplored with regards to high-throughput sequencing characterization of protist communities composition and spatial distribution of major taxonomic groups. This study contributes to fill this gap by investigating protist communities in relation to major water masses and oceanographic fronts characteristic of the SW Pacific waters east of New Zealand (Figure 1).

**Fig. 1.**
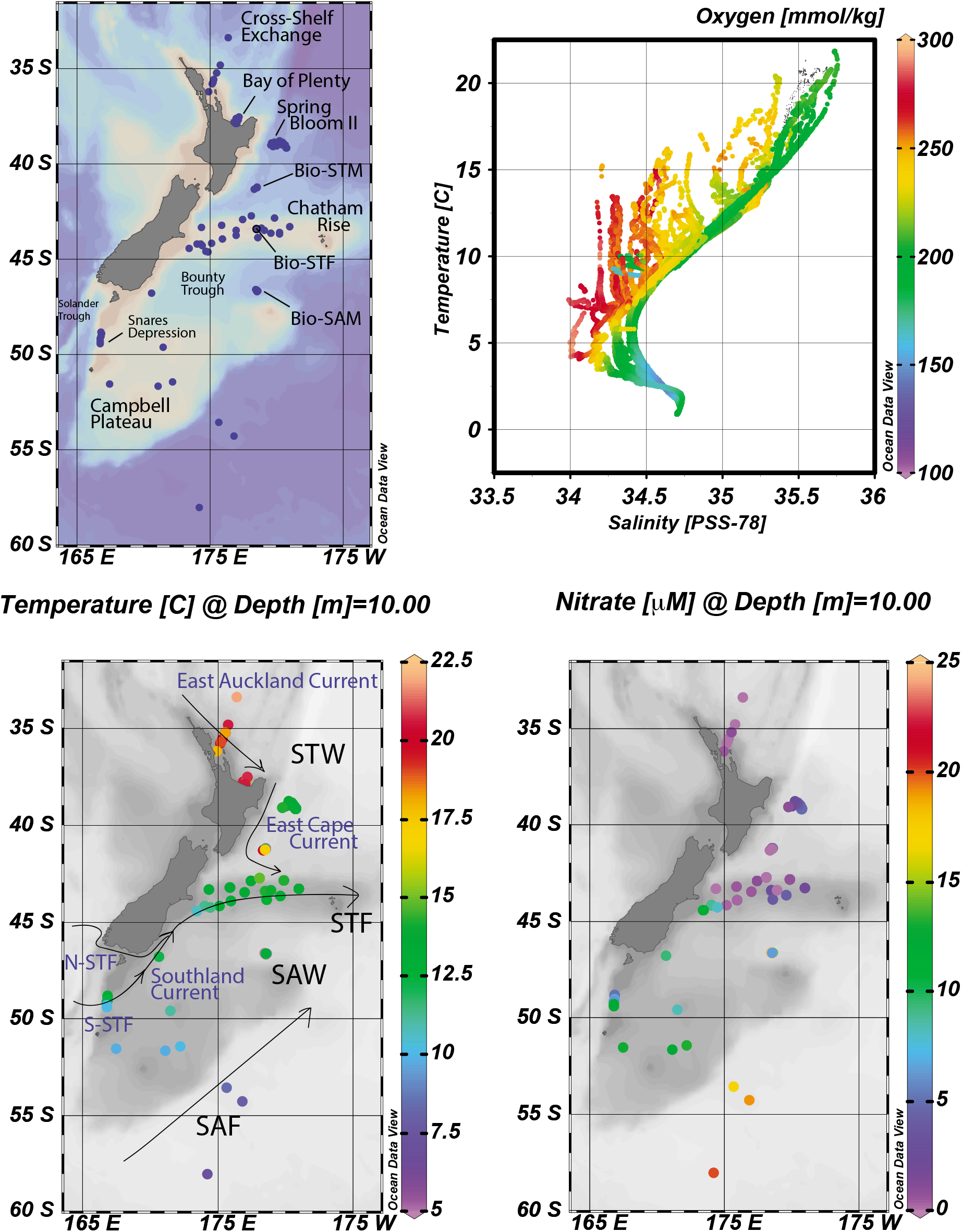
Study area. (A) Map of the study area with the sampling sites locations. (B) T-S diagram with oxygen concentration. Surface (C) temperature (C) and (D) nitrate concentration (D) at sampling sites in relation to major water masses and currents and fronts of the study region. North (N-STF) and South Subtropical Front (S-STF) adapted from Smith et al. 2013

New Zealand‘s continental mass interrupts the converging flows of the South Pacific subtropical gyre and the northward excursions of the Antarctic Circumpolar Current (ACC). The mixing of the warm and saltier subtropical water (STW) with the cold, relatively fresh subantarctic water (SAW) (Boyd et al. 1999) results in the genesis of oceanic fronts and semi-permanent eddies with distinctive signatures in water properties extending along the eastern margin off New Zealand (Fernandez et al. 2018). To the north, the East Auckland Current (EAUC) brings STW sourced partially by the Tasman Front (Sutton and Bowen 2014). At about 37°S the EAUC turns south to become the East Cape Current (Stanton et al. 1997) extending the STW inflow to the Chatham Rise where it separates from the coast to the east as part of the Subtropical Front (STF) (Deacon 1982). The STF is characterised by strong temperature gradients and a sharp salinity contrast that intensifies near the rise (Smith et al. 2013), up to 4°C and 0.7 practical salinity units respectively over 1° latitude in this region (Belkin and Gordon 1996). This transitional zone separating waters of subtropical origin from the subantarctic ones is known as the Subtropical Front Zone (SFTZ) (Deacon 1982) and it is bounded by the northern (N-STF) and southern (S-STF) branches of the STF. The STFZ can be up to 500 km wide in the Tasman Sea region before it gets constricted around the South Island of New Zealand where gradients set in motion, guided by the continental slope, the geostrophic flow associated with the S-STF branch and its coastal expression, the Southland Current (SC). The mean transport of the SC is about 8 Sv (1 Sv = 10^6^*m*^3^*s^−^*^1^) with 10% corresponding to STW and 90% to SAW (Sutton 2003). The SC advects this mix of STW and SAW northwards off the east coast and reaches south of the Chatham Rise through the Mernoo Gap and the Bounty Through. Further east and along the flanks of Campbell Plateau, the flows associated with the Subantarctic Front (SAF) carry the largest portion of SAW, about 50 Sv into the region south and east of the Chatham Rise (Bowen et al. 2014; Stanton and Morris 2004). Access of SAW onto the plateau from the east occurs through the bathymetric gaps, saddles and ridges where waters then become isolated from the neighbouring circulation and significantly contribute to the development of oceanographic and climatic processes such as subantarctic mode water formation (Forcén-Vázquez et al. 2021). Southeast of the Chatham Rise and away from the plateau the STFZ re-emerges as a 150 km wide band with the typical signatures of the STF-N and STF-S fronts (Sutton 2001).

STW and SAW have contrasting biogeochemical characteristics (Boyd et al. 1999; Bradford-Grieve et al. 1999; Chiswell et al. 2015; Heath 1985; Sherlock et al. 2007). North of Aotearoa New Zealand, STW is oligotrophic (low macro- and micronutrients) and phytoplankton production is considered to be limited by nitrogen (Zentara and Kamykowski 1981) with pervasive nitrogen-fixation by diazotrophs (Ellwood et al. 2018; Law et al. 2012). The STF is a dynamic region, characterized by strong temperature and salinity gradients (Sutton 2001) where high levels of vertical and lateral mixing of nitrogen-limited STW and macronutrient-rich SAW (Chiswell 2001), leads to regionally elevated annual net primary production (Murphy et al. 2001; Pinkerton et al. 2005). In SAW iron is the primary limiting nutrient for phytoplankton growth (Banse 1996; Boyd et al. 1999) although silicate and light can become limiting at times in SAW extending southeast of Aotearoa New Zealand which is considered high-nutrient, low-chlorophyll, low-silicate (HNLC-LSi) region (Boyd et al. 2010; Dugdale et al. 1995). These conditions are typically associated with SAW north of the Subantarctic Front (SAF), which is an area commonly referred to as the Subantarctic Zone (SAZ) (Trull et al. 2001) or the Subantarctic Water Ring (Longhurst 2007). In the SAZ, increasing availability of dissolved silica southwards shifts the Polar Frontal Zone extending between the SAF and the Polar Front to ‘standard’ Southern Ocean HNLC conditions (Rigual-Hernández et al. 2015).

Several studies have characterized microbial community composition in STW and SAW east of New Zealand, using microscopy (Chang and Gall 1998), pigment (Delizo et al. 2007) and flow-cytometry (Hall et al. 1999). These regional studies have focused mainly on the STF zone or coastal communities (Chang et al. 2003; Hall et al. 2006), while studies analyzing wider phytoplankton distribution across STW and SAW have targeted specific groups such as coccolithophores (Chang and Northcote 2016; Saavedra-Pellitero et al. 2014). More process-oriented studies have also provided partial information on phytoplankton composition in SAW and STW east of New Zealand (Chiswell et al. 2019; Ellwood et al. 2013; Peloquin et al. 2011). These studies have described the prevalence of larger cells and diatoms through winter and spring in the more productive waters of the STF, compared to STW and SAW. Diatom- and autotrophic flagellate-dominated communities have been reported in STW on the northern flank of the Chatham Rise during spring while dinoflagellates and small flagellates are documented as dominating the eukaryotic phytoplankton in SAW (Bradford-Grieve et al. 1997; Chang and Gall 1998). Diatom and coccolithophore species composition of sediment trap fluxes on the northern (STW-influenced) and southern (SAW-influenced) flanks of the Chatham Rise highlight the importance of these phytoplankton groups in the region (Wilks et al. 2021). However, there is surprisingly only little information available on the taxonomic composition of phytoplankton communities prevailing in open-ocean waters away from the STFZ over the Chatham Rise region (Chang and Northcote 2016; Chiswell et al. 2019; Peloquin et al. 2011; Twining et al. 2014). Further east of this region (170 °W), phytoplankton community composition from polar to equatorial waters have been characterized using pigment analysis (DiTullio et al. 2003) whereas a more recent study applied DNA metabarcoding analysis to investigate microbial diversity patterns in relation to physico-chemical gradients and oceanographic features (Raes et al. 2018). DNA metabarcoding has also been recently applied to investigate protist diversity changes across the Southland Current, a coastal expression of the STF that flows along the eastern margin of New Zealand’s South Island (Allen et al. 2020). However, the taxonomic composition of protistan communities associated with open-ocean water masses in the SW Pacific and across major oceanographic fronts that separate them is still lacking.

The aims of the present study are: 1) to characterize the diversity of protistan communities in STW and SAW east of New Zealand and across the STFZ that separates these water masses, and 2) to investigate the spatial distributional patterns of the main protistan taxonomic groups and species in relation to physical and chemical variability of the main water masses east of Aotearoa New Zealand. Specifically, we want to know how (dis-)similar are protist communities in the biogeochemically contrasting STW and SAW? What are the main environmental factors responsible for these differences? Which are the main taxonomic groups associated with each water mass and their environmental preferences? To do so we have applied DNA metabarcoding analysis (18S rRNA) to > 450 samples collected during 12 oceanographic voyages conducted over several years (2009-2017) and different seasons across STW and SAW east of New Zealand. This sequence data together with core physico-chemical and biological parameters (e.g. temperature, salinity, mixed-layer depth (MLD), macronutrients, total and size-fractionated chlorophyll a) provides the most comprehensive dataset of protistan plankton diversity in STW and SAW in the SW Pacific and contributes significantly towards building a robust baseline against which future changes in the region can be evaluated.

## 2. METHODS

### 2.1. Study area and sample collection

Seawater samples and data were collected during 12 research cruises conducted in SW Pacific waters east of Aotearoa New Zealand between 2009 and 2017 (Figure 1). The dataset covered 100 stations distributed between 54.3 and 33.4 °S with seawater samples (n = 482) collected over the 0-2000 m depth range and covering spring, summer and autumn periods (Table 1). The number of DNA samples from STW (n=269) were 2-fold higher than those from SAW (n=120) and STF (n=94) mainly due to the large number of samples from the Spring Bloom II voyage (TAN1212,(Chiswell et al. 2019)) (Table 1). The seasonality coverage was similar among the three different regions (STW, SAW, STF) but was biased against winter with most samples collected during spring, summer and autumn periods. (Table 1; Figure S1). Details about latitudinal and seasonal coverage of each water mass and the sample density distribution of analysed DNA samples, together with physico-chemical variables, are shown in Figure S2.

**Table. 1.** Summary of cruises from which samples were collected. Information includes the cruise identification code, start and end dates, the project or region, the water masses surveyed, latitudinal and seasonal coverage, number of stations and samples collected in each cruise.

| Cruise | Start | End | Project | Water mass | min Lat | max Lat | Season | N stations | N Samples |
| --- | --- | --- | --- | --- | --- | --- | --- | --- | --- |
| TAN0902 | 30-01-09 | 03-02-09 | BiophysMoorings | SAW // STF // STW | -46.62 | -41.23 | Summer | 3 | 28 |
| TAN0909 | 27-10-09 | 30-10-09 | BiophysMoorings | SAW // STF // STW | -46.64 | -41.2 | Spring | 3 | 32 |
| TAN1006 | 06-05-10 | 08-05-10 | BiophysMoorings | SAW // STF // STW | -46.64 | -41.19 | Autumn | 3 | 33 |
| TAN1103 | 19-02-11 | 21-02-11 | BiophysMoorings | SAW // STF // STW | -46.61 | -41.31 | Summer | 3 | 34 |
| TAN1113 | 29-09-11 | 01-10-11 | BiophysMoorings | SAW // STF // STW | -46.63 | -41.22 | Spring | 3 | 34 |
| TAN1203 | 17-02-12 | 05-03-12 | SOAP | STF | -44.61 | -43.48 | Summer | 10 | 10 |
| TAN1204 | 19-03-12 | 21-03-12 | BiophysMoorings | SAW // STF // STW | -46.64 | -41.26 | Autumn | 4 | 32 |
| TAN1212 | 19-09-12 | 05-10-12 | Spring Bloom II | STW | -37.87 | -37.51 | Spring | 19 | 105 |
| KAH1303 | 08-03-13 | 14-03-13 | Bay of Plenty | STW | -39.17 | -38.76 | Autumn | 12 | 37 |
| TAN1516 | 05-12-15 | 21-12-15 | Chatham Rise | STF | -44.42 | -42.72 | Summer | 20 | 39 |
| TAN1604 | 14-05-16 | 21-05-17 | Cross-shelf Exchange | STW | -36.18 | -33.38 | Autumn | 7 | 42 |
| TAN1702 | 18-03-17 | 29-03-17 | Campbell Plateau | SAW | -54.26 | -46.77 | Autumn | 13 | 52 |
| TAN1802 | 13-02-18 | 13-02-18 | SO-RossSea | SAW | -58.03 | -58.03 | Summer | 1 | 5 |

Samples were collected from 10 L Niskin bottles attached to a CTD rosette in association with a Seabird 9plus CTD, equipped with temperature, salinity, dissolved oxygen, fluorometer and beam transmissiometer sensors. K*_d_* PAR was estimated from chlorophyll a (Chl *a*) (Morel and Maritorena 2001). The euphotic zone depth (Zeu) was defined as the depth where downwelling PAR irradiance was 1% of incident irradiance (E_0_). The MLD was defined as the shallowest depth where density exceeded the 5 m value by 0.03 kg/m^3^ (Gardner et al. 1995). During the TAN1516 voyage samples were collected with a Niskin bottle deployed manually down to 10 m depth and from the R/V *Tangaroa* Underway Flow-Through System (TUFTS) system equipped with temperature, salinity, and fluorescence sensors. Seawater samples for nutrients, Chl *a* and DNA were sampled from the Niskin bottles using acid-washed silicone tubing and filtered through different types of filters for processing, as outlined below.

### 2.2. Nutrients, total and size-fractionated chlorophyll a

Samples for nutrients were filtered through 25 mm-diameter Whatman GF/F filters into clean 250 ml polyethylene bottles and frozen at −20 °C until analysis using an Astoria Pacific API 300 microsegmented flow analyzer (Astoria-Pacific, Clackamas, OR, United States) according to the colorimetric methods described in (Law et al. 2011).

For total Chl *a*, 250-400 mL seawater were filtered under low vacuum (<200 mm Hg) through 25 mm GF/F filters. These were folded and wrapped in aluminum foil or placed in Secol envelopes and stored at −80 °C or in liquid nitrogen until analysis. For size-fractionated Chl *a* (0.2-2 *µ*m, 2-20 *µ*m, >20 *µ*m) 400-500 mL were filtered sequentially through 47 mm polycarbonate filters by vacuum. Filters were folded and stored in 1.5 mL cryovials at −80 °C until analysis using 90% acetone extraction by spectrofluorometric techniques on a Varian Cary Eclipse fluorometer following method APHA 10200 H (Baird 2017)

### 2.3. DNA samples collection and extraction

Seawater samples of 1.5–5.0 L were filtered either through 0.22 *µ*m filters (47 mm-diameter polyether-sulfone, Pall-Gelman) using low vacuum or through 0.22 *µ*m Sterivex filter units (Millipore) using a peristaltic pump (Cole-Palmer). Disc filters were then folded and placed in cryovials and sterivex units were filled with RNAlater and flash-frozen in liquid nitrogen prior to storing at −80 °C. Disc filters were cut in two halves first and then into small pieces using a sterile blade. Each half was placed in separate tubes and lysed in parallel (2 h at 65 °C on a Boekel thermomixer set at 750 rpm) using the Nucleospin Plant kit Midi Kit (Macherey-Nagel, Duren, Germany). The 100 *µ*L of PL2 buffer recovered from each halved filter was then pooled together and the DNA extraction procedure continued with the Mini version of the Nucleospin Plant kit.

For Sterivex filters DNA was extracted using a Tris-buffered lysis solution containing EDTA, Triton X 100 and lysozyme (pH = 8.0) and the Qiagen DNeasy Blood & Tissue. Briefly, to collect cells that eventually detached from the filter surface, the RNAlater present in the filter unit was collected into a 2 mL Eppendorf tube using a syringe and then centrifuged (13,000 rpm, 10 min). The pellet was resuspended using 1 mL of the lysis solution and pipetted back into the original Sterivex. The cartridge was secured using Parafilm, put into a 50 mL falcon tube and incubated in a shaking incubator overnight (75 rpm, 37 °C). 1 mL of buffer Qiagen buffer AL and 40 *µ*L of proteinase K (20 mg/mL) was then added into the Sterivex. After securing the Sterivex, as described previously, the filter unit was incubated for 2 hours (75 rpm, 56 °C). Followed the incubations the lysate was recovered from the cartridge and DNA extraction and purification continued following manufacturer’s instructions in the Qiagen DNeasy Blood & Tissue.

### 2.4. PCR amplification, amplicon sequencing and sequence processing

The V4 region of the 18S rRNA gene was amplified using the eukaryotic primers V4F_illum (CCAGCAS-CYGCGGTAATTCC) and V4R illum (ACTTTCGTTCTTGATYRATGA) with Illumina overhang adapters (TCGTCGGCAGCGTCAGATGTGTATAAGAGACAG and GTCTCGTGGGCTCGGAGATGTGTATAA-GAGACAG) following procedures described in (Gutierrez-Rodriguez et al. 2019). PCR reactions were prepared in 50 *µ*L using 2x KAPA HifFi HotStart ReadyMix, (0.3 nM dNTP, 2.5 mM MgCl2), 0.5 *µ*M of each primer and of DNA template. The thermocycling profile included 95 °C/3 min, 10 cycles (98 °C/10s, 44 °C/20s, 72 °C/15s), 15 cycles (98 °C/10s, 62 °C/20s, 72 °C/15s) and 72 °C/7min).

Amplicon sequencing was conducted at the Genotoul GeT core facility (Toulouse, France) using an Illumina Miseq and a 2 x 250 cycles Miseq kit version 2. 482 samples were sequenced generating a total of 9166190 reads, with a median sequencing depth across all samples of 18954 reads per sample (range = 6539 – 36551). Obtained sequences were processed using the DADA2 pipeline (Callahan et al. 2017) following the procedure described by (Trefault et al. 2021). Taxonomic assignation was made against PR2 database version 4.12 (https://pr2-database.org/), yielding 16861 amplicon single variants (ASVs) assigned to protistan taxa. Details on the number of samples, reads and ASVs associated with each water mass are shown in (Table S1).

### 2.5. Pre-processing of ASV table and diversity analysis

We standardized the ASV table to the sequencing depth in each sample by normalizing the relative abundance to the mean number of sequences obtained across samples (median sequencing depth * (number of reads per ASV /total number reads per sample)). The relative contribution of specific groups in different water masses and regions were estimated from the sum of standardized reads across the samples considered.

Similarity analysis was undertaken using non-multidimensional scaling (nMDS) and ANOSIM using phyloseq (McMurdie and Holmes 2013) and vegan (Oksanen et al. 2019) R packages. Differences in species abundance across waters masses and regions was analysed using negative binomial generalized linear models coded in the DESeq2 package (Love et al. 2014). For analysis of higher taxonomic rank (division, class), distribution and their relation to environmental variables, the tax_glom function in phyloseq was used to agglomerate the previously standardized ASV table into the chosen taxonomic level.

### 2.6. Data accessibility

Physico-chemical, biological and geographic data including measurements of temperature, salinity, oxygen and transmissivity obtained with a Seabird 9plus; inorganic nutrients; total and size-fractionated chl *a*); and estimated mixed-layer and euphotic zone depth can be found PANGAEA repository archive data sets (submitted to PANGAEA). Raw sequence data are available on NCBI under accession number PRJNA756172 (). The quality filtered ASV table together with the taxonomy and metadata table used to build the phyloseq object are provided as csv tables. R scripts for data processing and figures can be found in https://github.com/agutierrez2001/Catalyst_Biogeography.

## 3. RESULTS

### 3.1. Physical, chemical and biological variability

STW was identified as those waters with surface salinity >35 psu (range = 35.1 - 35.6) (Figure 2) and included samples collected during the Cross-shelf Exchange II (TAN1604), the Bay of Plenty (KAH1303), the Spring Bloom II (TAN1212) voyages, and several cruises that visited the northern mooring site (Bio-STM) of the Biophysical Moorings long-term monitoring program (Nodder et al. 2016) (Figure 1, Table 1 and Figure S1). The Subtropical Front (STF) had salinity values ranging between 34.4 and 35.0 psu (Figure 2) and included samples collected on the Chatham Rise during TAN1516 (Fisheries Oceanography IV) and several Biophysical Mooring voyages as well as those collected in the STF upstream of the Chatham Rise, as it flows northwestward between the South Island and the Campbell Plateau (TAN1702) (Figure 1, Table 1, Figure S1). Typically SAW had salinity values <34.4 (Figure 2) and included samples collected at the southern Biophysical Mooring site located in the Bounty Trough (Bio-SAM), and at several stations on Campbell Plateau, and in the SAF south of the plateau (TAN1702, TAN1802) (Figure 1, Table 1, Figure S1).

**Fig. 2.**
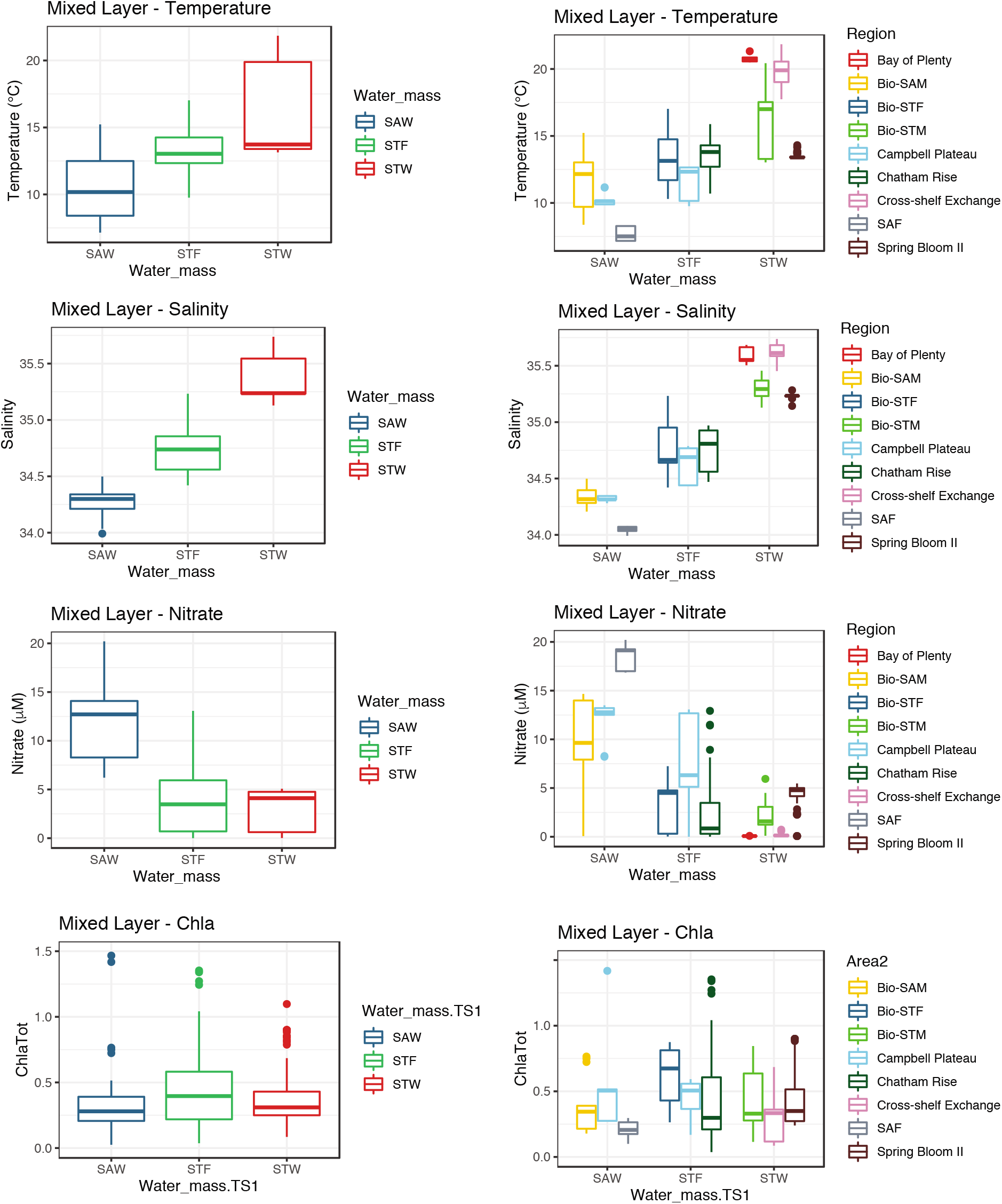
Surface mixed-layer physico-chemical variability. Box-plot representation of surface mixed-layer temperature and salinity, nitrate and chlorophyll a concentration in each water mass. Box-plots show the median, the first and third quartiles (lower and upper hinges) and the values within (line) and outside (dots) the *±*1.5 *∗ IQR* (IQR, interquartile range).

Sea-surface temperature was on average lowest in SAW (10.7 ± 2.4 °C, mean ± standard deviation, sd), intermediate in the STF (13.1 ± 1.7 °C) and highest in STW (16.1 ± 3.2 °C) (Figure 2). Temperature showed greater overlap among water masses and regions than surface salinity (Figure 2). STW sampled during the Spring Bloom II voyage, for instance, showed surface temperature consistently lower than 15 °C (12.5 - 14.5°C) (Figure 2). Nitrate concentrations were lowest in STW (2.98 ± 1.96 *µ*mol/L), intermediate and more variable in STF waters (4.28 ± 4.17 *µ*mol/L) and highest in SAW (12.17 ± 4.02 *µ*mol/L), consistent with HNLC conditions of these southern waters (Figure 1D).

Chl *a* concentration in the surface mixed-layer was on average higher in the STF (0.65 ± 0.65 *µ*g/L) compared to STW (0.38 ± 0.31 *µ*g/L) and SAW (0.37 ± 0.23 *µ*g/L) (ANOVA, F(2,220) = 14.2, p< 0.0001). Most of the DNA samples included in this study were taken from oligotrophic and mesotrophic waters (surface Chl *a* <0.5 *µ*g/L) with only a few collected from waters with Chl *a* concentrations >1.0 *µ*g/L. The smallest size-fraction (0.2 – 2.0-*µ*m Chl *a*) dominated the phytoplankton communities across all water masses but more so in STW (Chl *a* <2.0 *“’-*75 % of total Chl *a*) compared to SAW and STF (40-50%, Figure S3). The contribution of >20-*µ*m size-fraction to surface mixed layer Chl *a* was higher in SAW and STF, and although it remained on average relatively low (<15%), it occasionally reached > 50% levels (Figure S3).

A closer look revealed regional differences within each water mass (Figure 2). In SAW for instance, SAF surface waters were colder and fresher than those on the Campbell Plateau (TAN1702) and in the Bounty Trough (Bio-SAM site) (Figure 2). Surface nitrate concentrations were lower in the Bounty Through compared to Campbell Plateau and SAF, consistent with the southwards strengthening of HNLC conditions. *Chla_ML_* concentration was higher on Campbell Plateau (0.62 ± 0.48, mean ± sd) compared to surface waters in the Bounty Trough (0.33 ± 0.20) and the SAF (0.21 ± 0.07) (Figure 2), although differences were only significant between the Campbell Plateau and SAF to the south (one-way ANOVA, F(2,33) = 4.494, p = 0.019).

Within STW, surface temperature and salinity were highest in northernmost waters sampled during the Cross-shelf Exchange II voyage (TAN1604) and lowest in STW waters sampled during the Spring Bloom II voyage (TAN1212) conducted at the beginning of austral spring (September-October) (Figure 2). Temperature and salinity at the Biophysical Mooring subtropical site (Bio-STM) were intermediate on average and had a greater range that reflected the wider temporal variability covered by multiple visits conducted at different times of the year over several years (Table 1). Nitrate concentrations showed the opposite trend with highest values associated with the colder and deep-mixed waters, and lower values reflecting warmer and stratified waters of the Bay of Plenty and Cross-shelf Exchange II voyages (Figure 2).

Regional differences were also observed between the S-STF flowing north of the Campbell Plateau through the Snares Depression (Figure 1), which transported colder and fresher waters, and the STF further north flowing eastwards over the Chatham Rise (STF, Chatham Rise) (Figure 2). Nitrate concentrations tended to be higher in STF waters adjacent to Campbell Plateau than on the Chatham Rise reflecting the HNLC nature of the plateau. Relatively high nitrate concentrations (>10 *µ*mol/L) were also measured during summer (TAN1516) on the Chatham Rise at some stations located in the southwestern flank of the STF with colder (10.7 and 11.3 °C) and fresher (34.47 and 34.56) characteristics of surface mixed-layer waters indicating a SAW influence (Figure 1).

### 3.2. Alpha-diversity – Species richness

Species richness and the Shannon diversity index estimated in the euphotic zone were on average lower in the STF compared to SAW and STW (Figure 3A). Highest diversity in STW was observed in the northernmost waters surveyed during the Cross-shelf Exchange II voyage and at the Bio-STM site, which included samples collected during various cruises that spanned multiple years and seasons. Protist species richness during the Spring Bloom II voyage was substantially lower than these other STW locations (Figure 3B). In SAW diversity was lower on Campbell Plateau compared to open ocean regions adjacent to flows of the SAF (Figure 3B) located to the north and south of the plateau, respectively (Figure 1). Within the STF, diversity was also lower in upstream waters of the S-STF located further south (46-49 °S) than in the STFZ in the Chatham Rise region (43-45 °S) (Figure 1, Figure 3). Here, higher diversity values were associated with the Bio-STF site visited multiple times in different seasons during the Biophysical Mooring program (2009-12)(Table 1) than during the summer TAN1516 voyage, which covered the Chatham Rise region more extensively (Figure 1).

**Fig. 3.**
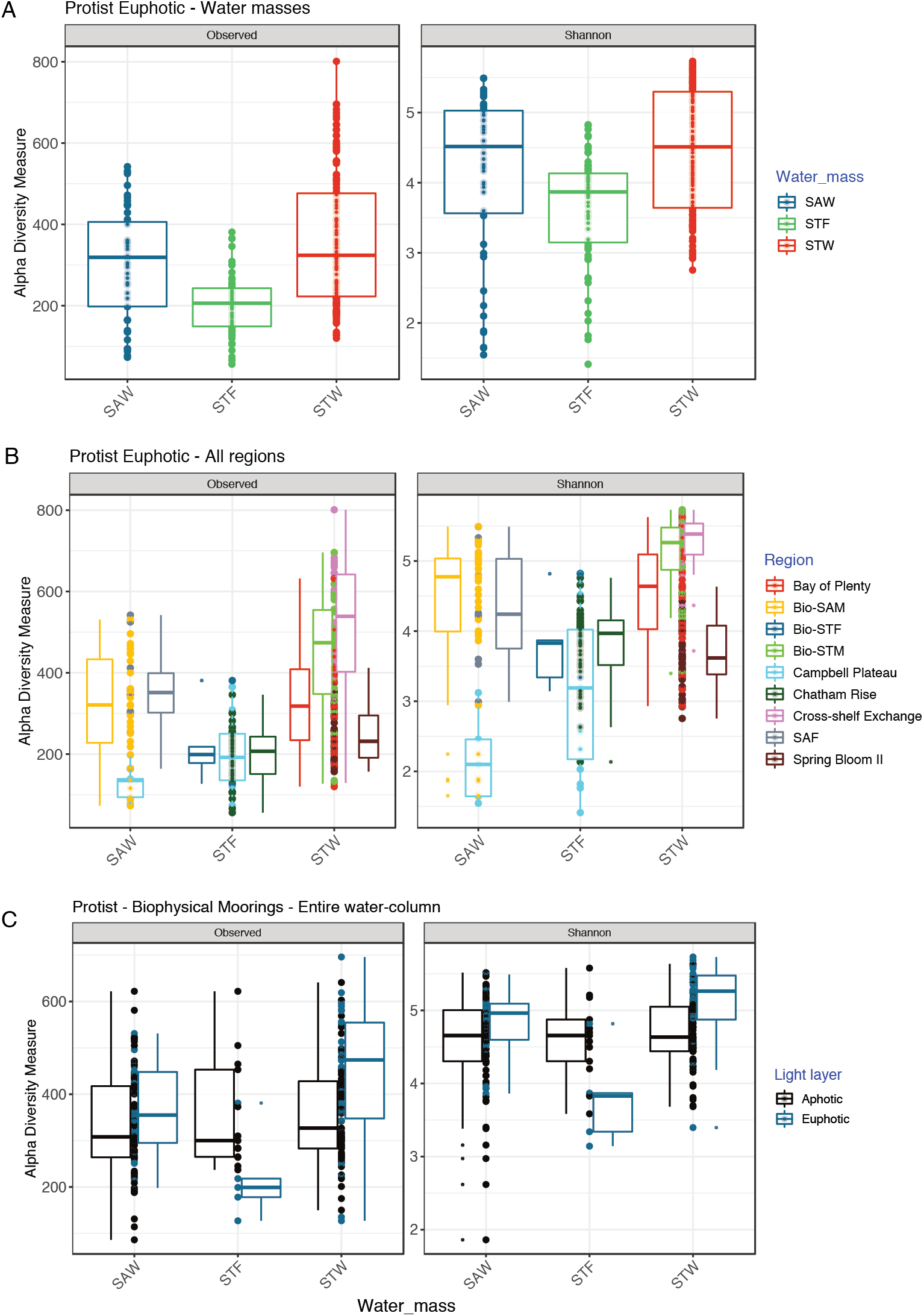
Species richness and diversity index estimated for (A) the euphotic zone of each water masses (SAW, STF, and STW); (B) the euphotic zone of each regions and (C) the aphotic zone of the Biophysical Mooring programme sites SAF (Subantarctic Front) and Campbell Plateau correspond to the same voyage TAN1702 (April 2017).

Species diversity in the euphotic zone tended to decrease with latitude (model I linear regression, F(1,236) = 25.6, R2 = 0.10, p-value < 0.0001), although differences in mean diversity values were observed among water masses and regions with similar latitudes (Figure S4). Similar relationships were observed between diversity and temperature (model I linear regression, F(1,208) = 59.1, *R*^2^ = 0.20, p-value < 0.0001) with regional differences modulating this trend (Figure S5). Within STW for instance, species richness in the euphotic zone was higher in oligotrophic waters and decreased with increasing Chl *a*, being generally higher in warmer waters (Figure S6). Samples from the STF presented lower species richness in the euphotic zone compared to SAW and STW across the entire range of Chl *a* and nitrate concentrations (Figure S6).

Diversity patterns in the aphotic zone were investigated on the Biophysical Moorings dataset only (Bio-STM, -STF and -SAM, n = 113). Diversity in the aphotic layer of the STW and SAW sites was lower than in the sunlit layers (Figure 3C). In the STF, however, this was higher in the aphotic compared to the euphotic layer, which resulted in similar estimates of species richness in the aphotic zone of the different water masses unlike the diversity pattern observed for the euphotic zone (Figure 3).

### 3.3. Beta-diversity of protistan communities

To explore the similarities between protistan communities in different geographic samples we performed principal component analysis on ASV abundance. The first analysis with all samples yielded two main clusters corresponding to samples from the euphotic and aphotic zone (Figure 4A). A second analysis focused on the euphotic zone cluster samples (n = 239) according to different water masses although certain overlaps also occurred, particularly between the STF and SAW samples (Figure 4B). Different regions tended to cluster separately as well, with STW samples from the Spring Bloom II voyage forming a separate cluster from that of the Cross-shelf Exchange and Bay of Plenty cruises (Figure 4B).

**Fig. 4.**
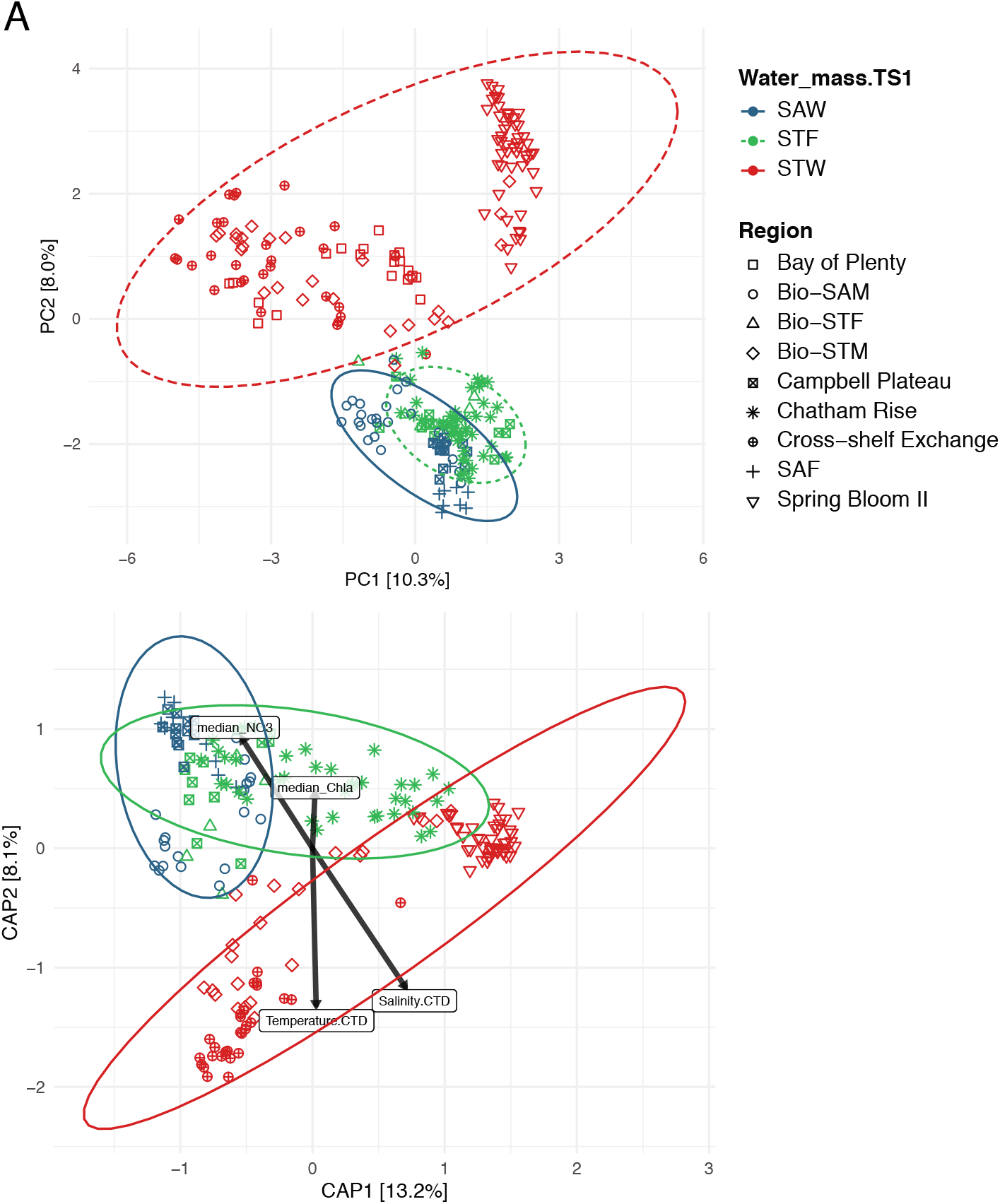
Figure 4. (A) Principal component analysis based on ASV composition of euphotic samples only color coded by water masses and shapes for regions/voyages (n=240). (B) Biplot of redundancy analysis (RDA) computed at species (ASV) level in the euphotic zone for which T, Sal, Nitrate and Chl *a* measurements were available. Arrows indicate the sign and strength of the correlation between community composition an environmental variables that were significant in PERMANOVA analysis (n=197) samples from Chatham Rise TAN1516 lack CTD and MLD data

To investigate the influence of different oceanographic drivers on the community composition we performed PERMANOVA analysis with continuous variables of physical (T, Sal), chemical (NO*^−^*_3_ concentration - euphotic zone median NO_3_^−^) and biological (Chl *a* concentration - water-column median) parameters. The analysis was conducted on a subset of samples (n=182) for which these measurements were available (STW=34, STF=53, SAW=95). All variables yielded statistical significance (p<0.001) with salinity explaining most of the variability (F.Model = 34.4, R^2^ = 0.13, P<0.001) followed by temperature (R^2^=0.08) and nitrate (R^2^=0.06) with chla concentration explaining a minor fraction of this variability (R^2^=0.02) (Figure S7) (Table S2). Overall this set of variables left 69% of the variability unexplained. The second PERMANOVA analysis included categorical variables (e.g. Water mass and region) showed that while Water mass explained 16% of the variability (F. Model = 28.0, P<0.001) – similar to that explained by salinity – the region explained an additional 27% of the variability (F.Model = 15.4, P<0.001) and up to 51% of the variability together with physico-chemical and biological continuous variables included in the first PERMANOVA analysis (Table 2).

**Table. 2.** Summary of PERMANOVA analysis including the Water-mass and Region as categorical variables in addition to the continuous environmental variables. Temperature and salinity represent the surface values. Nitrate (N0*^−^*_3_) and chlorophyll a (Chl *a*) median concentration calculated for samples within the euphotic zone. Analysis was conducted with the Adonis function of the vegan R package

| Variable | Df | SumsOfSqs | MeanSqs | F.Model | R<sup>2</sup> | Pr(>F) |
| --- | --- | --- | --- | --- | --- | --- |
| Water-mass | 2 | 9.321 | 4.6606 | 27.6669 | 0.16054 | 0.001 |
| Region | 6 | 15.558 | 2.593 | 15.3932 | 0.26796 | 0.001 |
| Temperature | 1 | 1.459 | 1.4588 | 8.6597 | 0.02512 | 0.001 |
| Salinity | 1 | 1.05 | 1.0503 | 6.2351 | 0.01809 | 0.001 |
| median NO<sub>3</sub> | 1 | 1.076 | 1.0763 | 6.3893 | 0.01854 | 0.001 |
| median Chl *a* | 1 | 1.128 | 1.1283 | 6.6977 | 0.01943 | 0.001 |
| Residuals | 169 | 28.469 | 0.1685 |  | 0.49032 | 0.001 |
| Total | 181 | 58.062 |  |  | 1 |  |

### 3.4. Division and class level taxonomic composition

Dinoflagellate reads (syndiniales excluded) dominated the sequencing datasets from samples taken in the euphotic zone (34% of total - sequencing depth-normalized – reads), with Chlorophyta accounting for 27%. Ochrophyta, constituted mainly by *Bacillariophyta* (7%) and *Pelagophyceae*(3%)) and *haptophytes* belonging to *Prymnesiophyceae* class(5%) were the other most important phytoplankton divisions, while Stramenopiles_X (mainly through Marine Stramenopiles, MASTs, 4%) Radiolaria (4%), and Ciliophora (3%) contributed most among the heterotrophic protist. Although such groups were consistently dominant, their relative contributions and particularly, their composition at class (Figure 5 and Figure 6) and finer taxonomic resolution (see subsection 3.5.) varied between water masses.

**Fig. 5.**
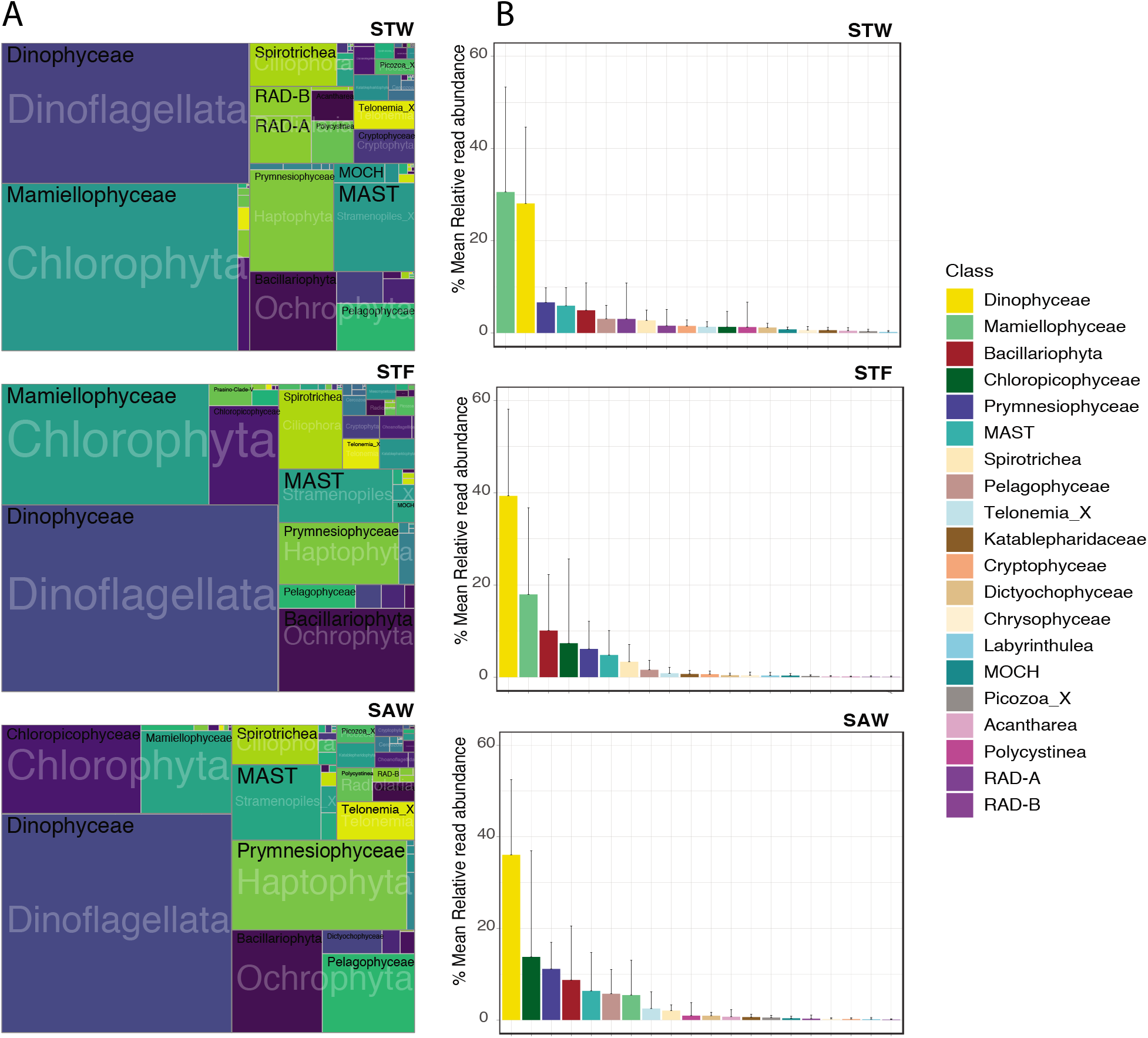
Protist community composition at division/class level (syndiniales excluded) in the euphotic zone of STW, STF and SAW. A) The area of each taxonomic group in the treemap represents the read abundances affiliated to each group standardized to the median sequencing depth across samples [median sum otus * (otu reads / sum (otu reads)]. B) Barplots represent the mean relative read abundance of most abundant classes across different water masses (error bars are the standard deviation of the mean).

**Fig. 6.**
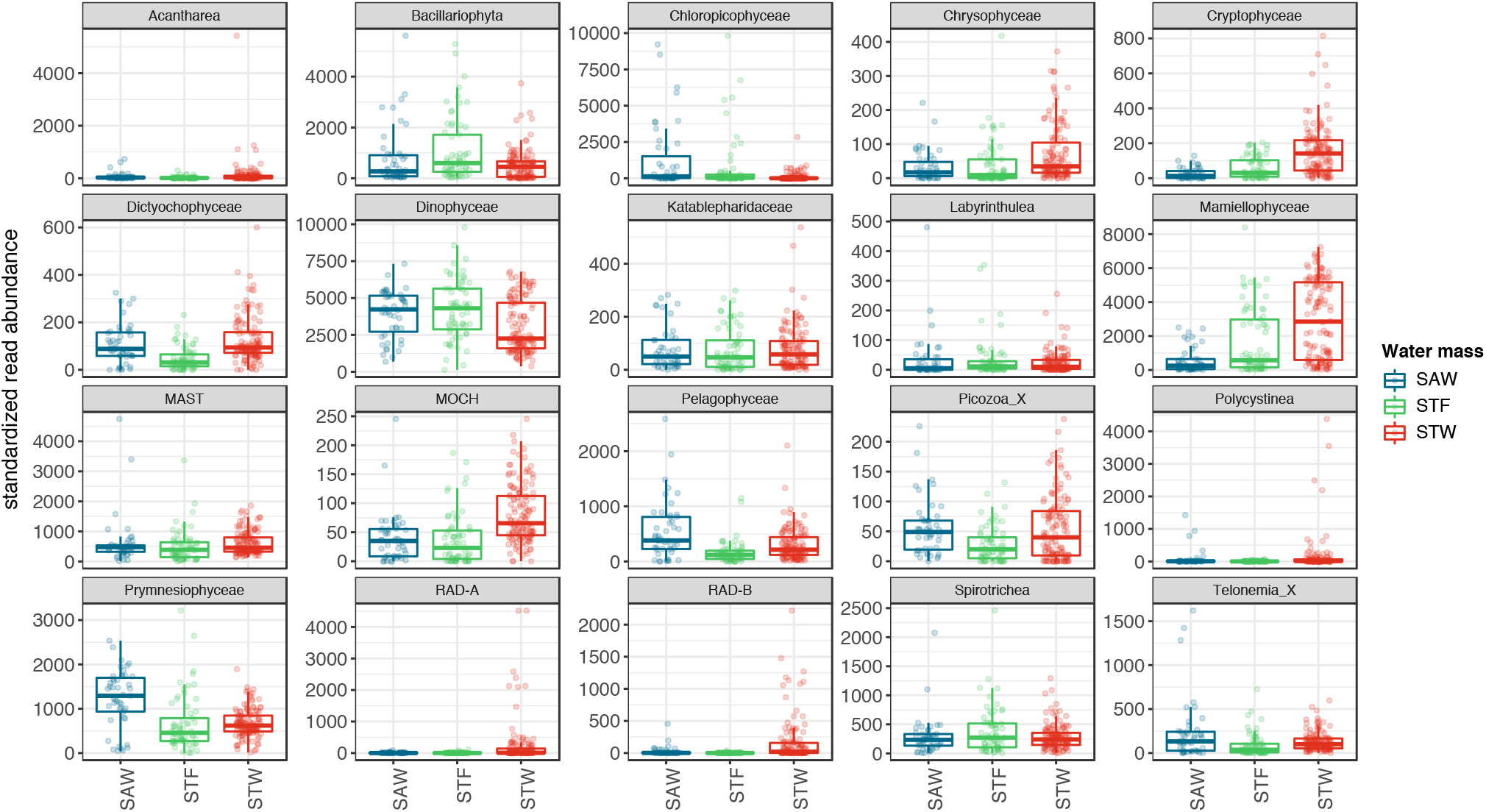
Box-plots showing standardized read abundance in subtropical, subantarctic and subtropical front of the twentieth most abundant protist classes in the euphotic zone. Box-plots show the median, the first and third quartiles (lower and upper hinges) and the values within the *±*1.5 *∗ IQR* (IQR, interquartile range) (line). Points represent values of single samples.

For STW samples for instance, where dinoflagellates and Chlorophyta co-dominated the overall dataset (30% of sequences each), followed by Ochrophyta (10%) with Haptophyta, Stramenopiles_X and Radiolaria accounting for a lower percentage of metabarcodes (7% each). *Mamiellophyceae* accounted for the vast majority of reads affiliated to Chlorophyta (>95%) while *Chloropicophyceae* represented only a minor fraction of sequences affiliated to this division (Figure 5). *Prymnesiophyceae* was the most abundant class of Haptophyta (6%). *Bacillariophyta* (5%) and *Pelagophyceae* (3% each) classes dominated over *Dictyochophyceae* (1%) and *Chrysophyceae (1%)* among Ochrophyta, while *Cryptophycea* (Cryptophyta, 2%) accounted for a relatively larger percentage. The class Among Heterotrophic groups MASTs (Stramenopile_X, 5%), RAD-A (Radiolaria, 3%) and ciliates of the class *Spirotrichea* contributed most to the STW metabarcoding data (Figure 5 and Figure 6).

The SAW sequencing dataset was clearly dominated by dinoflagellates (37% total reads), followed by Chlorophyta (18%), Ochrophyta (15%) and Haptophyta (12%) with a more even share of the total number of reads compared to STW (Figure 5). At class level, *Chloropicophyceae* (14%), was clearly the most abundant group of green algae followed by *Prymnesiophyceae* (12%), *Bacillariophyta* (8%) and *Pelagophyceae* (6%) phytoplankton classes (Figure 5). The heterotrophic component in the SAW metabar-coding data was dominated by MASTs (6%) followed by ciliates (3%) while the relative contribution of Radiolaria in the euphotic zone was minor (<0.5%) and mainly attributed to Polycystinea rather than RAD-A &-B classes (Figure 5 and Figure 6).

The STF protist metabarcoding data presented intermediate characteristics between STW and SAW (Figure 5). As in SAW, dinoflagellates (39%) dominated the data although the contribution of Chlorophyta (26%) was on average higher and closer to levels found in the STW metabarcode dataset. Accordingly, *Mamiellophyceae* was the second most abundant class of green algae (17%), although the contributions of *Chloropicophyceae* (10%), and *Pyramimonadophyceae* (1.5%) were also substantial. The division Ochrophyta (12%) accounted for a similar fraction of phytoplankton reads as in STW and SAW, but in STF the phytoplankton community was clearly dominated by *Bacillariophyta* (10%) with minor contributions from *Pelagophyceae* (2%) and *Dictyochophyceae* (0.5%) classes. The heterotrophic component was dominated by MASTs (5%) and ciliates (4%) while the contribution of Radiolaria (<0.2%) was on average lower than in STW and SAW datasets (Figure 5 and Figure 6).

To investigate protist community composition below the euphotic zone we used the Bio-physical moorings dataset that covered systematically the entire water column at the Bio-STM and Bio-SAM (0-3100 and 0-2800 m, respectively) sites and the Bio-STF site on the Chatham Rise crest (0-350 m)(n = 113 samples). Overall, Radiolaria (Polycystinea) dominated the aphotic metabarcoding data (52% of protist reads) followed by dinoflagellates (27%). Heterotrophic groups such as MASTs (3%) and ciliates (3%) contributing substantially less (Figure 7 and Figure S8). Phytoplankton groups such as diatoms, green algae, and prymnesiophytes contributed at similar levels (3% each), likely reflecting mixing layers extending below the euphotic zone. The relative contribution of Radiolaria was similar in SAW (57%) and STW (50%) samples but much lower in the STF samples (35%) where together with dinoflagellates (36%) they co-dominated protistan reads below the euphotic zone (Figure 7 and Figure S8). Among Radiolaria, Polycystinea class, mainly through the Spumellarida order, was most abundant (25-35% of total reads) although Acantharea (10%), RAD-B (8%) and to a lesser extent RAD-A (1%) classes also contributed to the dominance of this group (Figure 7 and Figure S8).

**Fig. 7.**
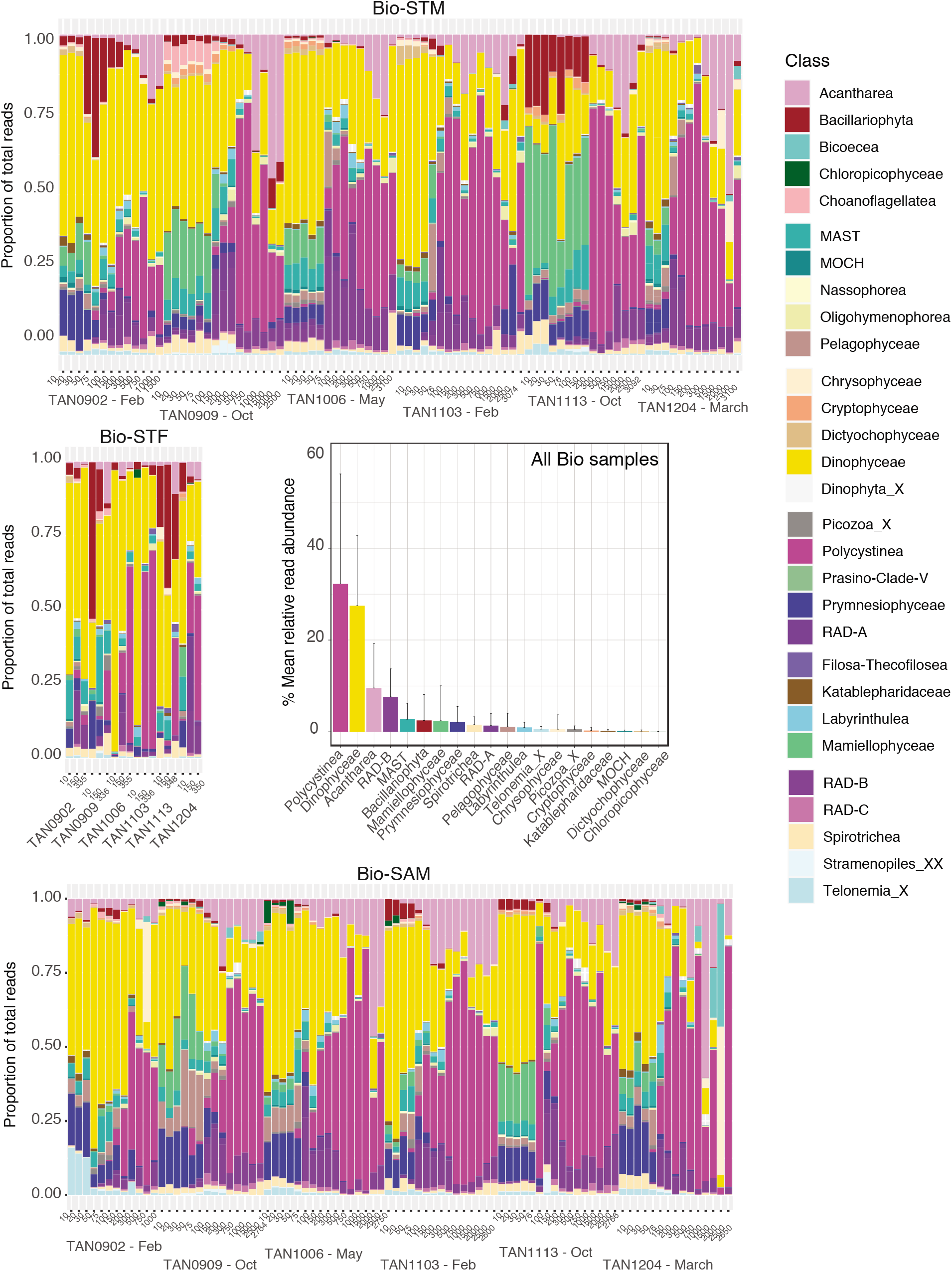
Relative read abundance of main protistan classes in samples collected throughout the water column (0-2000 m) during multiple voyages to the Biophysical Moorings program sites in STW (Bio-STM), STF (Bio-STF) and SAW (Bio-SAM) and mean contribution for the whole sampling program (n = 113)(error bars as in Figure 4.

Vertically, the dominance of Radiolaria became more prominent in the mesopelagic and bathypelagic samples where they often represented >75% of total reads (Figure 7). Polycystinea class tended to dominate across all sample depths, with increasing abundance at depths >300 m. Sequences affiliated with Acantharea and RAD-B showed similar abundance but followed opposite vertical distributional trends, with RAD-B being more abundant in shallower samples (100-500 m) while Acantharea abundance increased with depth and peaked in bathypelagic samples (>1000 m) (Figure 7).

### 3.5. Genus and species community composition

Species composition also varied among the metabarcodes from the physically and biogeochemically distinct water masses (Figure 8, Figure 9). In STW, the green algae (Chlorophyta) was dominated by the species *Ostreococcus lucimarinus* followed by *Bathycoccus prasinos*. These *Mamiellophyceae* species together with *Micromonas commoda* and other *Micromonas* species (*M. bravo* I, II, and *M. pusilla*) accounted for the majority of the sequences affiliated to green algae in the euphotic zone of STW (Figure 8, Figure S9 and Figure S10). Reads affiliated to several dinoflagellate species such as *Gymnodinium* sp., *Heterocapsa rotundata* and *Gyrodinium spp.* were among the most abundant in STW dataset. *Gymnodinium* sp., and *H. rotundata* were more abundant at the Bio-STM whereas *Gyrodinium helveticum* was prevalent in STW of the EAUC surveyed during the Cross-shelf Exchange voyage (Figure S10). Diatom ASVs identified as Polar-centric Mediophyceae_X sp. and *Minidiscus trioculatus* were the most abundant diatoms reads in STW metabarcodes, particularly in the Spring Bloom II voyage. Among ASVs assigned to pelagophytes, an unidentified Pelagophyceae_XXX_sp. (ASV_0058) and *Pelagomonas calceolata* ASVs were the most abundant ones (Figure 8, Figure S10 and Figure S9). Among the class *Prymnesiophyceae*, *Phaeocystis globosa* (ASV_0065) was the most abundant ASV (Figure 8) with several ASVs belonging to *Gephyrocapsa oceanica* and *Chrysochromulina* spp. mainly contributing to the overall dominance of the latter genus within the class (Figure S9, Figure S10).

**Fig. 8.**
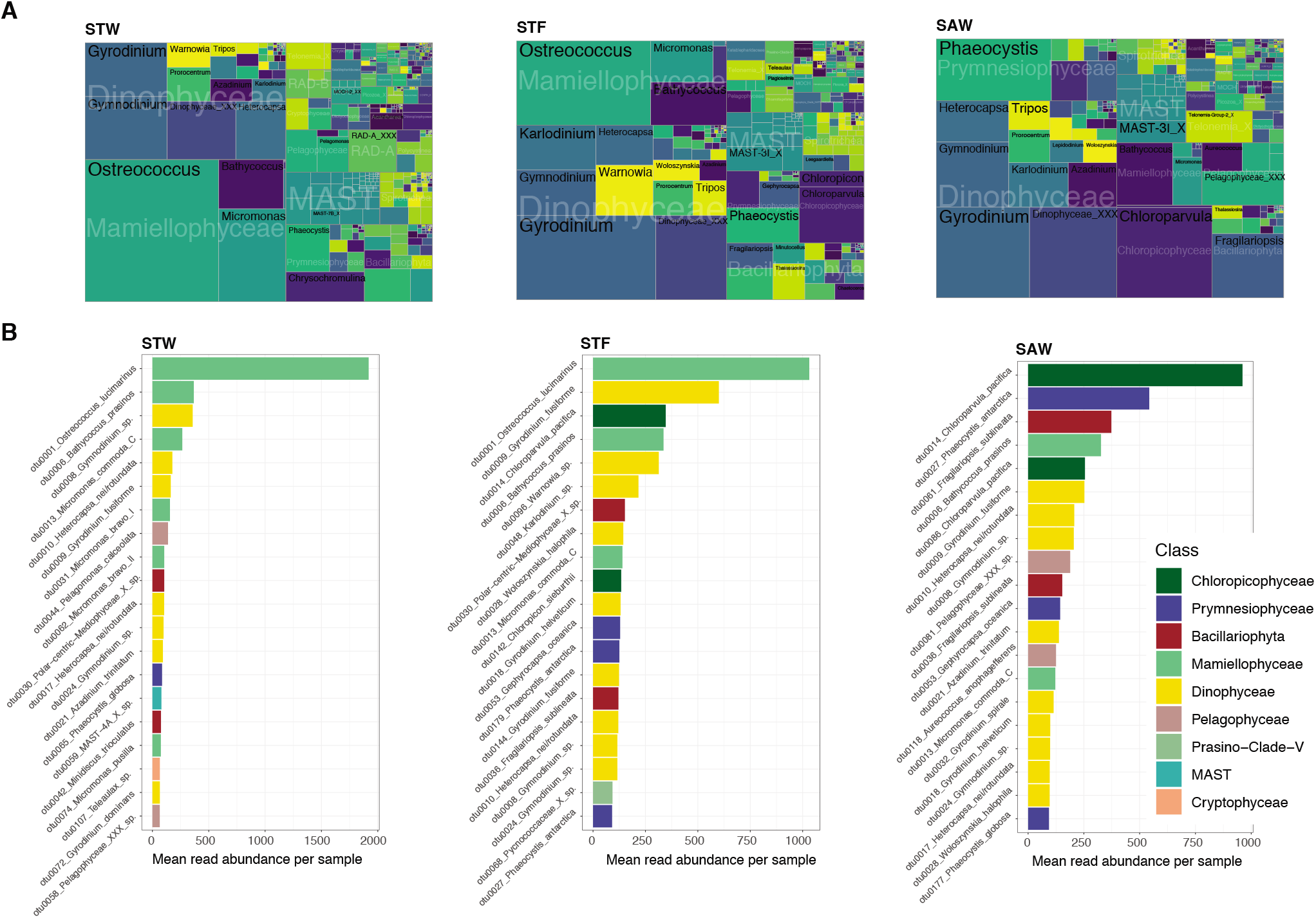
Water mass genus and species abundance (A) Treemaps showing the community composition at class/genus level in the euphotic zone of the STW, STF and SAW. The area of taxonomic group is proportional to the read abundances affiliated to each group standardized to the median sequencing depth across samples [median sum otus * (otu reads / sum (otu reads)]. (B) Mean standardized read abundance of most abundant ASVs and assigned species, color coded for their class affiliation, in the euphotic zone of the different water masses.

**Fig. 9.**
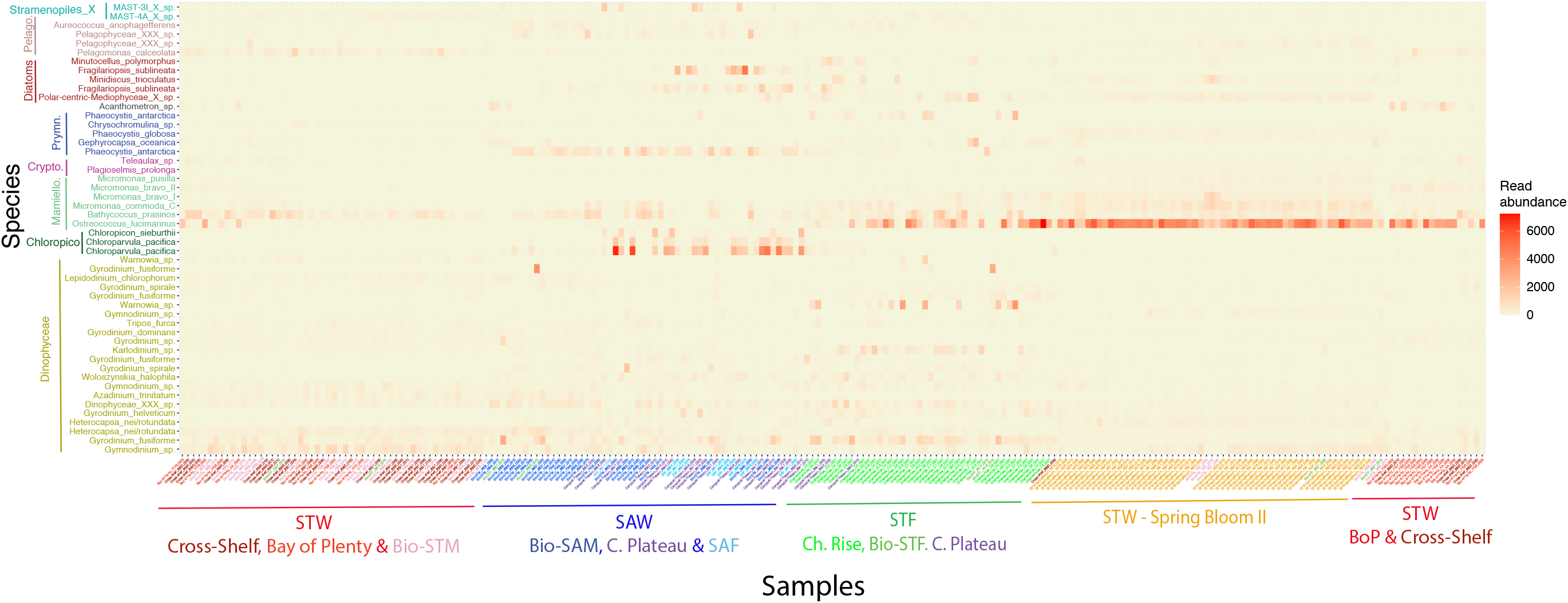
Heatmap showing the standardized read abundance of the 50 most abundant species (Y-axis) across samples collected in the euphotic zone (X-axis). Samples were clustered using nMDS and Jaccard distance and sample labels color coded according to the water mass and region they were collected from. Species were organized and color coded by class affiliation.

Among the STF metabarcodes, the Mamiellophyceae *O. lucimarinus* was also the most abundant species overall, although the relative contribution of *Chloroparvula pacifica* (asv_0014, *Chloropico-phyceae*) increased substantially compared to STW (Figure 8, Figure S9). It is worth noting the high abundance of sequences affiliated to *Chloropicon sierburthii* in addition to *Chloroparvula pacifica*, which contributed to the overall increase in the relative abundance of *Chloropicophyceae* in the STF dataset (Figure 8, Figure S9). Reads assigned to the heterotrophic dinoflagellate *Gyrodinium fusiforme* was the second most abundant in STF samples, with other ASVs affiliated with *Warnowia* sp. and *Karlodinium* sp. appearing among the most abundant dinoflagellate species (Figure 8, Figure 9 and Figure S9). The higher relative abundance of Bacillariophyta observed in STF was mainly driven by ASVs affiliated to unidentified Polar-centric Mediophyceae species and *Fragilariopsis sublineata* (Figure 8 and Figure S9) which tended to be more abundant in the STFZ of the Chatham Rise region (TAN1516) and the S-STF flowing north of C. Plateau (TAN1702), respectively (Figure S10). The identification of *F. sublineata* should be taken with caution the V4 region of the two sequences included in PR2 is 100% similar to the annotated sequence of *Fragilariopsis Kerguelensis* present in PR2 making impossible to unambiguously assign that ASVs to one of this two species. To reflect this ambiguity the ASVs assigned to *F. sublineata* in PR2 are referred as *F. sublineata/kerguelensis* througout the text (see discussion). Among *Prymnesio-phyceae*, *Phaeocystis* spp. was the dominant genus but most reads in this case belonged to *P. antarctica* instead of *P. globosa* (Figure 8, Figure S9). ASVs assigned to the prymnesiophyte *G. oceanica* also increased substantially in STF compared to STW datasets. *Pelagophyceae* in the STF metabarcodes was dominated by *Aureococcus* and *Pelagococcus* spp. (Figure S9 by although the class relative contribution was relatively low (Figure 5). The relative abundance of *Cryptophyceae* and *Dictyophyceae* remained minor overall (<2%) (Figure 5), but both groups showed changes in their specific composition across the water masses metabarcode datasets. Among *Cryptophyceae*, *Plagioselmis prolonga* and *Teleaulax gracilis* increased substantially from STW to STF samples while the relative contribution of sequences assigned to *Teleaulax* sp. decreased. Changes within *Dictyophyceae* were less substantial but showed an increase in the relative abundance of *Dictyocha speculum* and *Pseudochattonella farcimen* from STW to STF metabarcodes (Figure S9).

Among SAW samples, *Chloropicophyceae* ASV_0014 assigned to *Chloroparvula pacifica* was the most abundant taxa, with other ASVs of this species (e.g., asv_0086) contributing also to the dominance of this class over Mamiellophyceae in SAW (Figure 8, Figure 9, Figure S9). Among Mamiellophyceae, reads from *B. prasinos* became the most abundant followed by *M. commoda* while the contribution of *O. lucimarinus* in SAW samples was minor (Figure 8 and Figure S9). The increase in the relative abundance of Prymnesiophyceae reads observed in SAW samples was driven mainly by *Phaeocystis* spp., with *P. antarctica* emerging as the second most abundant species in SAW metabarcodes (Figure 8). Other *Phaeocystis* species (*P. globosa*, *P. cordata* and *Phaeocystis* spp.) contributed also to the dominance of this genus within prymnesiophytes (Figure S9 and Figure S10). Among Bacillariophyta, ASVs assigned to *F. sublineata/kerguelensis* (ASVs 0061 and 0036) were the dominant in SAW samples (Figure 8 and Figure 9) while *Thallasiossira* sp., Polar-centric Mediophyceae sp. and other diatom genus contributed substantially less (Figure S9 and Figure S10). Increasing abundance of pelagophyte reads in SAW (Figure 5) was mainly due to *Pelagophyceae*_XXX_sp (ASV_0081), the same ASV that dominated in STW samples, and to *Aureococcus anophagefferens* which were among the most abundant taxa in SAW metabarcodes (Figure 8 and Figure S9).

To investigate how the relative abundance of the species identified varied between the water masses, we ran a differential gene expression analysis based on the negative binomial distribution (DESeq) using water mass as a categorical variable. For the euphotic zone, this yielded 70 and 35 ASVs out of 3984 ASVs that were significantly more or less abundant (p-value < 0.01) in STW compared to SAW datasets, respectively (Figure S11A). Species that showed greatest differences (>10 log2-fold changes) in their relative abundance were not necessarily among the most abundant species in each water mass. Among the species associated with STW, we found the larger changes in relative abundance in the diatoms ASVs assigned to Polar-centric Mediophyceae, *Minutocellus polymorphus*, and *Minidiscus trioculatus*; the prasinophytes *O. lucimarinus* and several *Micromonas species*; the prymnesiophytes *Phaeocystis globosa* and *Chrysochromulina* sp., and dinoflagellates *Warnovia* sp., *H. rotundata* and number of other species (Figure S11). Among species with preferences for SAW we found several diatom species including *F. sublineata* and *Cylindrotheca closterium*, the prasinophyte *B. prasinos*, the pelagophytes *Pelagococcus* sp. and *A. anophagefferens*, and several dinoflagellates including the heterotrophic species *G. fusiforme* and *G. spirale* and mixoplankton *Karlodinium veneficum* (Figure S11).

Different ASVs affiliated to the same species often showed distinct preference for STW and SAW. Most abundant *Chloroparvula pacifica* ASVs (e.g. asv_0086 and asv_0014, Figure 8) were associated with SAW, while less abundant ASVs (e.g. asv_0532 and asv_0336) were more associated with STW (Figure S11A). Similarly, most abundant ASVs of *P. antarctica* (asv_0011, asv_0027) showed preference for SAW (10-30-log2 fold negative change) while much less abundant asv_0218 showed greater affinity for STW. This intraspecific variability was observed within unidentified species (e.g. Dinophyceae_XXX_- sp. and Pelagophyceae_XXX_sp.) which included ASVs with opposite affinities for STW and SAW (Figure S11A).

DESeq analysis between the STF and STW also depicted specific differences as observed between STW and SAW, with some additional diatoms ASVs assigned to *Thalassiosira* sp. and *Pseudo-nitzschia delicatissima* enriched in the STF dataset (Figure S11B). In most cases, the distinctive species pattern observed between the STF and either STW or SAW metabarcodes coincided with those identified from the comparison between STW and SAW described above (Figure S11B). For instance, ASVs of Polar-centric Mediophyceae species found in higher abundance in STW compared to SAW, were also associated preferentially with STF compared to SAW while *F. sublineata*, and *C. closterium* diatoms ASVs, that were enriched in SAW compared to STW, were also preferentially associated with the STF rather than STW (Figure S11B). Only few ASVs, such as those assigned to the prasinophyte *Chloropicon sieburthii* and dinoflagellate *Gonyaulax* sp., were distinctively associated with STF waters (Figure S11C).

The most abundant reads found below the euphotic zone belong to the polycystinean order Spumellarida Group I family (Figure S8). Several ASVs contributed to the dominance of Spumellarida Group I family, but different ASVs dominated in each water mass dataset (Figure S12 and (Figure S13). ASV0023 and ASV0039 affiliated with the Group I family of Spumellarida dominated in SAW, while ASV0064 and ASV0085 were most abundant in STF (Figure S13). Conversely, most abundant radiolarian species in STW were unidentified species assigned to the acantharean order Chauncanthida (Figure S13). Several less abundant Radiolaria ASVs showed significantly different abundances across the aphotic depth layers of different water masses suggesting differences in their ecological preferences (Figure S14). Reads assigned to known photosynthetic species also showed differential abundance below the euphotic zone in the different water masses datasets. Most notably the higher abundance of reads affiliated to the *Mamiellophyceae O. lucimarinus* and the *Prymnesiophyceae P. globose* in STW compared to SAW yielded significant differences in their abundance below the euphotic zone as well. A pattern observed also for less prominent phytoplankton species like the diatom *Asterionellopsis glacialis* or *Thalassiosira rotula* that despite being less abundant in the euphotic zone yielded significant differences in their abundance below the euphotic zone.

## 4. DISCUSSION

The taxonomic composition of protistan communities in SAW and STW in the SW Pacific and across the main frontal zones of the region (STF, SAF) has has not been extensively characterized. Previous studies have typically focused on phytoplankton communities and their variability across SA and ST waters flanking the STFZ over the Chatham Rise (Chang and Gall 1998; Delizo et al. 2007; Hall et al. 1999) but broader biogeographic studies in the SW Pacific are scarce (DiTullio et al. 2003). By using DNA metabarcoding, we provide a comprehensive taxonomic characterization of the protistan communities associated with STW and SAW at southern temperate latitudes. Although samples included in this study were collected in different seasons, the seasonal coverage was similar for STW and SAW (Table 1 and Figure S1), allowing a meaningful comparison among the microbial communities associated with these water masses throughout an average year.

### 4.1. Protistan community structure in STW and SAW of the southwest Pacific

Species richness decreased latitudinally and with temperature (Figure S4 and Figure S5) as expected from global diversity patterns observed and modelled for marine bacterial and phytoplankton communities (Barton et al. 2010; Fuhrman et al. 2008; Ibarbalz et al. 2019) as well as from previous DNA-based reports in the SW Pacific region (Raes et al. 2018). Consistent with this trend, species richness was higher in warmer STW than in SAW while lowest diversity was associated with the STF (Figure 3).

The relative low diversity observed in the STF could be related to the increased phytoplankton biomass and productivity typically associated with the STF across the annual cycle (Murphy et al. 2001; Pinkerton et al. 2005) and the dominance of fewer ‘bloom-forming’ species in this highly productive zone (Chang and Gall 1998). The fact that species richness within STW was lowest during the more productive spring bloom conditions (TAN1212) is consistent with the view that more productive waters such as found in the STF, during the spring bloom, and locally on the Campbell Plateau (Gutiérrez-Rodríguez et al. 2020) tend to decrease protistan diversity. However, the lower diversity associated with STF, relative to STW and SAW, was systematically observed across the different levels of nitrate and Chl *a* concentrations encompassed in this study (Figure S6) suggesting that other factors may contribute to this pattern. Interestingly, the lower diversity in the STF relative to STW and SAW was buffered below the euphotic zone (Figure 3C). Similarly, the temperature-driven latitudinal pattern described globally for epipelagic plankton also disappeared below the euphotic zone (Ibarbalz et al. 2019). The decoupling between the diversity patterns in the sunlit and dark ocean suggested by these results are somewhat contrary to the connectivity between the epi- and bathypelagic zones as inferred by the high correspondence of bacterial communities and processes between these realms (Mestre et al. 2018; Ruiz-Gonzalez et al. 2020). The reasons for these differences are unclear and highlight the need of further studying ecological processes that shape microbial diversity throughout the entire water column.

Despite the regional and seasonal variability encompassed within both STW and SAW (Table 1, Figure 2) we observed systematic differences in the taxonomic composition associated with these water masses (Figure 4). Such water-mass specificity has been previously observed for the prokaryotic (Agogué et al. 2011; Galand et al. 2010; Seymour et al. 2012; Techtmann et al. 2015) and eukaryotic components (Hamilton et al. 2008; Raes et al. 2018) of microbial communities across different oceans. Among environmental drivers, salinity rather than temperature or nitrate concentration was the physico-chemical variable that explained best the compositional (dis-)similarities among euphotic samples (Figure 4, Figure S7). Bray-Curtis dissimilarity indices of surface bacterial communities across the Southland Current was also strongly correlated with salinity (Baltar et al. 2016). These results support the view that STW and SAW east of New Zealand are better conceptualized as bioregions or provinces rather than habitats sensu (sensu Martiny et al. 2006), where (phyto)plankton communities reflect oceanographic processes and history in addition to contemporary physico-chemical conditions.

Samples from the STF itself were also distinguished from those in SAW and STW based on their taxonomic composition, although they showed a greater overlap (Figure 4) that reflected the active mixing and transition nature of such frontal zones. This overlap was particularly evident between samples collected at the Bio-STF and Bio-SAM sites located on the Chatham Rise and its subantarctic flank, respectively, and between the S-STF and SAW over the Campbell Plateau, which suggests a stronger physical connectivity and ecological affinity of the STF with SAW than STW. Similarly, the horizontal mixing and phytoplankton community size structure in the STF zone has been reported to be more tightly coupled across SAW-influenced than STW-influenced water types (Safi et al., submitted). Nevertheless, the distinct protistan communities observed in STW and SAW, and to a lesser degree in the STF, highlights the role of oceanographic features such as the STF as boundaries that influence the diversity of oceanic microbial communities in large oceanic provinces (Baltar et al. 2016; Raes et al. 2018).

### 4.2. Taxonomic composition of phytoplankton community

Our results showed the overall dominance of dinoflagellates and Chlorophyta across all water masses, followed by Bacillariophyta, *Prymnesiophyceae* and *Pelagophyceae* (Figure 5). Yet consistent differences in the relative contribution of these taxonomic groups between water masses emerged at class and species taxonomic classification levels (Figure 6, Figure 8, Figure S9). Analysis of intra-specific diversity revealed differences in the distribution of ASVs of the same species suggesting the presence of different ecotypes in some cases (e.g. *Chloroparvula pacifica*, *P. antarctica*) and current taxonomic gaps within certain groups that remain to be characterized (e.g. *Pelagophyceae*_XXX; Dinophyceae_XXX)(Figure S11). Below we discuss the distributional patterns of major taxonomic groups, highlighting different taxonomic ranks to shed some light on their ecological preferences.

#### 4.3.3. Chlorophyta (Green algae)

The relative contribution of the two main green algae classes, *Mamiellophyceae* and *Chloropicophyceae*, showed opposite distribution patterns (Figure 5). *Mamiellophyceae* was the most abundant class in STW and constituted the bulk of green algae that dominated these waters samples, while its relative abundance decreased across the STF to reach lowest levels in SAW (Figure 6). Picoplanktonic algae *O. lucimarinus* was clearly the most abundant species in STW and STF (Figure 8, Figure 9) in agreement with a previous metabarcoding analysis conducted across the Southland Current (Allen et al. 2020) where this species abundance peaked in neritic STW inshore of the main current core and decrased towards SAW end of the coast-offshore sampling transect (Allen et al. 2020). *O. lucimarinus* was also among the most abundant species of *Mamiellophyceae* in a 18S rRNA metabarcoding survey conducted on coastal waters globally (Tragin and Vaulot 2019).

The dominance of picoplanktonic *Mamiellophyceae* in STW is consistent with the greater contribution of <2 *µ*m Chl *a* (80%,) observed in this water mass (Figure S4). It is worth noting that the highest abundance of this group was observed during the onset of the Spring bloom samples (TAN1212, Figure S10) when *Mamiellophyceae* accounted for 40-75% of 18S rRNA reads, while diatom contributions remained around 10% over the 3-weeks of sampling (Figure S15). Among the multiple surveys of the Bio-STM site, *Mamiellophyceae* contribution tended to be highest during early spring coinciding with the onset of the spring bloom (Figure 7). *Mamiellophyceae* and *O. lucimarinus* were the most abundant phytoplankton class and species in the STF samples along the Chatham Rise (TAN1516, Figure S15 and Fig, S10) particularly on the STW influenced northern flank of the rise (Figure S16). The abundance and presence of some prasinophyte classes, including *Mamiellophyceae*, have often been assessed from their diagnostic pigment prasinoxanthin. Quantitative application of pigment-based approaches showed that prasinophytes dominated the community in the STFZ and its subtropical flank across the Chatham Rise (170 °E) (Delizo et al. 2007) and further east (170°W) (DiTullio et al. 2003), in agreement with our DNA based results. In the Indian sector of the SO, a latitudinal study also found the highest contribution of prasinophytes associated with the STF (Iida and Odate 2014). Broader application of pigment approaches have revealed that prasinophytes can contribute substantially to vernal blooms at temperature latitudes (Bustillos-Guzman et al. 1995; Gutiérrez-Rodríguez et al. 2011; Latasa et al. 2010; Nunes et al. 2018). High abundance of several species of prasinophytes including *Ostreococcus* spp. and *Micromonas* spp. have been recently reported during the onset of the North Atlantic spring bloom from 16S rRNA amplicon sequencing analysis (Bolaños et al. 2020) and at more temperate latitudes of the Eastern North Atlantic using 18S rRNA metabarcoding (Joglar et al. 2021). The deep mixing layers (>100 m) during the New Zealand STW spring bloom II voyage (TAN1212) (Chiswell et al. 2019), where prasinophytes dominated, supports the idea that this picophytoplankton group thrives under high-nutrient, high-mixing conditions playing an important role in the development of spring blooms, characteristic of temperate latitudes. Overall, our results highlight the wide ecological breadth of *Mamiellophyceae* and certain species like *O. lucimarinus* which tend to dominate across a wide range of physical, chemical and trophic conditions encountered within STW.

*Chloropicophyceae* (previously defined as Prasinophytes clade VII, Lopes dos Santos et al. 2017a) showed an opposite trend to *Mamiellophyceae*, with the highest relative abundance associated with SAW samples (Figure 5 and Figure 6). Culture representatives of *Chloropicophyceae* and 18S rRNA sequences have been obtained from tropical and subtropical latitudes of the north and south Pacific (Lopes dos Santos et al. 2017a,b; Tragin and Vaulot 2018) but to our knowledge this is the first report of their presence and high abundance in subantarctic waters. The majority of *Chloropicophyceae* reads were assigned to a reference sequence corresponding to *Chloroparvula pacifica* and included several ASVs one of which (ASV0014) was the most abundant protist ASV found in SAW (Figure 8). *Chloropicophyceae* has been suggested as the dominant group of green algae in meso- and oligotrophic oceanic waters in contrast with the preference of *Mamiellophyceae* for more nutrient-rich coastal environments (Lopes dos Santos et al. 2017b; Shi et al. 2009; Tragin and Vaulot 2018). In this study, *Chloropicophyceae* were most abundant in SAW which are considered HNLC, suggesting that the preference of this group for meso-/oligotrophic conditions reported for typically macronutrient limited waters could also encompassed iron limited HNLC conditions. Furthermore, field experiments have suggested that phytoplankton growth in HNLC regions in the subarctic Pacific and the Southern Ocean is co-limited by B vitamins and iron micronutrients (Bertrand et al. 2007; Koch et al. 2011; Panzeca et al. 2006). Genome analysis of one species of *Chloropicophyceae* (*Chloropicon primus*) indicates that this group might be able to synthesize thiamine, in contrast to *Mamiellophyceae*, which depends on exogenous vitamin B1 or related precursors supplied by B1-synthesizing marine bacteria or other algae (Lemieux et al. 2019; Paerl et al. 2015). The potentially significant ecological role of B1 and B-vitamins, in general, in regulating and shaping the taxonomic composition of phytoplankton communities, is rarely considered and is still not well understood (Sañudo-Wilhelmy et al. 2014) but could contribute to explain the contrasting distribution patterns of both classes.

Although Chloropicophyceae abundance occasionally peaked in the Bounty Trough (Bio-SAM samples) the high relative contribution of this group in SAW was mainly due to the high abundance systematically observed on Campbell Plateau and the S-STF flowing north of the plateau (Figure 1, Figure S15, Figure S16). In the S-STF, *Chloropicon sieburthii* made a substantial contribution in addition to the more dominant *Chloroparvula pacifica* (Figure S10). Whether this regional preference was linked to the bathymetric and hydrographic characteristics of the plateau (Forcén-Vázquez et al. 2021; Neil et al. 2004), the natural iron fertilization hypothesized for the region (Banse and English 1997; Gutiérrez-Rodríguez et al. 2020) or a combination of these and other aspects cannot be concluded from our study. Moreover, an ASV belonging to this genus was also found to contribute substantially to protistan communities in coastal waters of the California Current Ecosystem (Gutierrez-Rodriguez et al. 2019), highlighting the need of further studies to better understand the ecological drivers beyond coastal-oceanic trophic gradients responsible for the water mass preferences of such phytoplankton groups and species.

#### 4.3.4. Dinophyceae (Dinoflagellates)

Dinoflagellates relative abundance tended to be higher in SAW and the STF compared to STW metabar-codes (Figure 5, Figure 6), consistent with previous microscopy-based observations (Chang and Gall 1998). ASVs affiliated to *Gyrodinium* genus and particularly *G. fusiforme* were identified as the most abundant species in agreement with a previous study in the Southland Current where DNA barcodes of *Karlodinium* and *Gymnodinium*, *Gyrodinium helveticum* and *G. spirale* were also retrieved and among the most abundant species (Allen et al. 2020). Many *Gyrodinium* species prey on bacteria and algae (Hansen 1992; Jang et al. 2019; Jeong et al. 2008). They can constitute an important component of microzooplankton biomass in coastal and oceanic environments (Jeong et al. 2010; Sherr and Sherr 2007) including high latitude waters (Archer et al. 1996; Olson and Strom 2002; Strom et al. 2001) where they have shown the capability of cropping down iron-stimulated diatom blooms (Saito et al. 2006). While species of *Gyrodinium* were prevalent across all water masses in our study, their abundance was higher in more productive STF waters (Figure 8, Figure S9), where higher Chl *a* concentrations was accompanied by increased abundance of diatoms and larger phytoplankton cells, confirming their pivotal importance in pelagic foodwebs as the link between primary producers and metazoan zooplankton (Zeldis and Décima 2020).

#### 4.3.5. Bacillariophyta (Diatoms)

Diatoms tended to be more abundant in the STF metabarcodes compared to STW and SAW although relatively high contributions (>30%) were at times attained in all water masses (Figure 6, Figure 7). Most abundant ASVs in STF waters were identified as polar-centric Mediophyceae_X sp., *Thalassiosira* sp. and *Fragilariopsis sublineata* (Figure 8, Figure S9), in the later case mainly due to their abundance in the southern STF flowing next to C. Plateau (Figure S10). Several diatom species including *F. sublin-eata/kerguelensis*, *F. cylindrus*, *Chaetocerus peruvianus*, and *Cylindrotheca closterium* were significantly more abundant in the STF compared to STW, but not compared to SAW (Figure S11) supporting the greater resemblance of diatoms assemblage between STF and SAW.

In SAW, the most abundant diatom ASVs were identified as closely related to *F. sublineata/kerguelensis* (Figure 8) consistent with the preference of *Fragiolariopsis* species for SAW inferred from microscopy analysis (Chang and Gall 1998). *F. sublineata* has been reported to dominate in sea ice algal biomass and for being well adapted to low light conditions (McMinn et al. 2010); however it is seldom reported among the dominant species in Southern Ocean surface waters, where other species like *F. curta* and *F. kerguelensis* tend to dominate (Mohan et al. 2011; Olguín and Alder 2011; Quéguiner et al. 1997) supporting the identification of ASVs assigned to *F. sublineata/F. kerguelensis* in this study to *F. kerguelensis*. However, the taxonomic assignation of the most abundant *F. sublineata/F. kerguelensis* ASVs (e.g., asv_0036,asv_- 0061) were only closely related to referenced sequences in PR2 (the bootstrap value at species level assignation of ASVs <50%) and their sequence showed 5 to 7 mismatches with the annotated sequences (Figure S17) which highlights the intraspecific diversity of the *Fragilariopsis* spp. The low silicate characteristics of SAW east of New Zealand (Dugdale et al. 1995) is likely a key factor responsible for the southward increase of heavily silicied diatoms like *Fragilariopsis* spp. which tended to be lowest showed in the Bio-SAM, intermediate on C. Plateau and highest in southern most waters of the SAF (Figure S10) in a way consistent with their tendency to dominate south of the SAF (Assmy et al. 2013; Pinkernell and Beszteri 2014). Furthermore, the shift in the relative abundance of the dominant ASVs assigned to *F. sublineata/kerguelensis* (ASV0036 and ASV0061) observed between subantarctic waters north (Bio-SAM, C. Plateau) and south of the SAF (SAF) (Figure S11, Figure S17) suggests potential differences in their silicate requirements for phytoplankton growth.

In STW, in addition to unidentified Polar-centric Mediophyceae_X sp., other small species such as *Minidiscus trioculatus* and *Minutocellus polymorphus* were identified among the most abundant diatoms (Figure 8, Figure S9), consistent with the dominance of these small diatom taxa in neritic-modified STW of the upstream Southland Current (Allen et al. 2020). While most common genera reported in STW (and STF) by microscopy analysis (e.g. *Thalasiosira* spp., *Chaetoceros* spp., *Guinardia* spp.) were also detected by DNA metabarcoding, the small nano-sized species revealed as numerically dominant by DNA approaches can be overlooked by microscopy (Chang and Gall 1998). Diatoms are generally conceptualized as the microplankton group associated with new production and high export potential (Legendre and Lefevre 1995; Uitz et al. 2006; Vidussi et al. 2001). However, there are increasing evidence supporting the importance of small nano- and even pico-sized diatoms in both coastal and oceanic systems (Arsenieff et al. 2020; Buck et al. 2008; Hernández-Ruiz et al. 2018; Lomas et al. 2009). Our results showing the dominance of *M. trioculatus* and *M. polymorphus* in STW particularly during the more productive conditions of the Spring Bloom II and the STF over the Chatham Rise (Figure 9, Figure S10) further support the important role played by small diatoms in driving open-ocean phytoplankton production (Leblanc et al. 2018).

#### 4.3.6. Pelagophyceae

Pelagophyceae showed the opposite trend compared to diatoms and have their lowest abundance associated with the STF zone (Figure 6). Their abundance and relative contribution increased towards SAW (Figure 5, Figure 6 in agreement with pigment-based dominance of *Pelagophyceae* in the SA waters east of C. Plateau (DiTullio et al. 2003). The relative abundance of this class and *Pelagomonas* species also decreased following natural or experimental iron addition experiments in HNLC waters of the SO (Irion et al. 2020; Thiele et al. 2014). These observations are consistent with the physiological advantage in iron uptake of pelagophytes over other small eukaryotic phytoplankton groups (Timmermans et al. 2005) and indicates a competitive advantage for pelagophytes under oligotrophic conditions. Vertically, the relative contribution of this class increased with depth (Figure 7, Figure S16) in agreement with their preference for deeper layers (Cabello et al. 2016; Gall et al. 2008; Latasa et al. 2017) and their physiological adaptation to low light and high nutrient environments (Dimier et al. 2009; Dupont et al. 2015). This vertical segregation was evident in both STW and SAW samples despite the different specific composition observed in each water mass (Figure S9) with *Pelagomonas calceolata* (ASV0044) and unidentified pelagophyte (Pelagophyceae_XXX.sp, ASV0081) being the most abundant species in STW and SAW, respectively (Figure 8)(Figure S9). *P. calceolata* is a widespread species (Andersen et al. 1996; Moon-van der Staay et al. 2001). Whether the ubiquity of this species is bound to high genetic diversity or physiological versatility is not clear. In our study, several ASVs were assigned to *P. calceolata* and while the most abundant one showed preference for STW, other less abundant ASVs were significantly more abundant in SAW (Figure S11). Similarly, we found different water mass preferences among ASVs assigned to unidentified pelagophytes, with some preferentially associated with STW or SAW but interestingly none with STF (Figure S11). While these observations suggest that different ASVs may reflect ecologically relevant different units (Rodríguez et al. 2005) they also highlight the importance of culture isolations and species characterization to better determine the diversity of pelagophyte assemblages.

#### 4.3.7. Prymnesiophyceae

Prymnesiophyceae were prevalent across all water mass metabarcodes (Figure 5) but tended to be more abundant in SAW (Figure 6). Overall, their relative contribution to eukaryotic phytoplankton assemblages was lower than depicted by pigment-based analyses of open ocean microbial communities (Andersen et al. 1996). The prevalence of 19’hexanoyloxyfucoxanthin pigment marker in oceanic waters and the application of quantitative methods (e.g. CHEMTAX) have shown that *Prymnesiophyceae* represents between 20-50 % of the phytoplankton community in oceanic waters (Andersen et al. 1996; DiTullio et al. 2003; Latasa et al. 2005; Liu et al. 2009; Swan et al. 2016). Such dominance has been also depicted by improved genomic approaches that revealed the extremely high genetic and functional diversity of non-calcifying prymnesiophytes (Cuvelier et al. 2010; Liu et al. 2009). In our study, non-calcifying species, mainly assigned to *Phaeocystis* spp. and *Chrysochromulina* spp., dominated the group (Figure S9) in agreement with DNA-based studies in the SW Pacific region (Sow et al. 2020; Wolf et al. 2014). The abundance and relative contribution of *Phaeocystis* spp. was lowest in STW, intermediate in STF and peaked in SAW while *Chrysochromulina* spp. followed the opposite trend with higher contributions associated with STW (Figure S9). The dominance of *Phaeocystis* spp. in SAW was mainly driven by *P. antarctica* (Figure 8), corresponding with the prominence of this group in the Southern Ocean (Verity et al. 2007) and observed decreasing abundance observed from SAW towards STW of the SW Pacific region during austral autumn-Winter (Sow et al. 2020). These results contrast with the similar spatial distribution *P. antarctica* spp. metabarcodes observed between contrasting conditions on and off the Kerguelen Plateau (Irion et al. 2020). Strains assigned to *P. globosa* and *P. cordata* were also detected in all water masses although they tended to be more prevalent and abundant in STW compared to SAW in the New Zealand region (Sow et al. 2020). Coccolithophores are an important component of phytoplankton communities in the Southern Ocean region extending from the STF to the Polar Front known as the Great Calcite Belt (Balch et al. 2016; Chang and Northcote 2016). *Gephyrocapsa oceanica* was the most prevalent and abundant coccolithophore species found in our study. ASVs assigned ot this species were found across all water masses but tended to be most abundant in the STF (Figure S9; Figure S10) in agreement with previous microscopy-based studies in this region of the SW Pacific (Rigual-Hernández et al. 2020; Saavedra-Pellitero et al. 2014). *Emiliania huxleyii*, which generally dominate coccolithophore assemblages in this region (Chang and Northcote 2016; Saavedra-Pellitero et al. 2014), and in the Southern Ocean (Balch et al. 2016; Holligan et al. 2010) showed very low abundances across the different water masses surveyed in this study (data not shown).

#### 4.3.8. Cryptophyceae

In our datasets, the contribution of *Cryptophyceae* metabarcodes was relatively low on average (<3%) but showed increasing abundance from SAW to STW where they represented up to 10% of protistan reads in the euphotic zone (Figure 6). The genus and species composition of this group also differ substantially between STW and SAW in our dataset (Figure S9). Similar water-mass preference was depicted by quantitative pigment analysis in the same STFZ region over the Chatham Rise, where cryptophytes contribution in STW (47-63% chl *a*) was higher than in SAW (6% chl *a*) in one of the two consecutive springs surveyed (Delizo et al. 2007). Cryptophytes are an ubiquitous phytoplankton group with widespread distribution from coastal to open oceanic systems and from tropical to polar latitudes (Buma et al. 1992; Nunes et al. 2019; Piwosz et al. 2013). They have been reported to form blooms in coastal embayments (Jeong et al. 2013; Johnson et al. 2013) and coastal Antarctic waters favoured by low salinity conditions (Moline et al. 2004; Nunes et al. 2019; Schofield et al. 2017). The higher contributions we observed in STW relative to SAW, however, argues against the direct influence of salinity on cryptophytes at least in open-ocean waters. Cryptophytes have been also observed to respond positively to iron fertilization in HNLC waters of the North Pacific (Sato et al. 2009; Suzuki et al. 2009) suggesting that their lower abundance in SAW in our study could be related to iron-limited conditions characteristic of the subantarctic region. The contribution of cryptophytes in STW was highest during the open-ocean spring bloom (Spring Bloom II-TAN1212 voyage) and in shelf-slope stations of the EAUC current (Cross-shelf Exchange-TAN1604 voyage) consistent with their preference for more nutrient-rich conditions (Carreto et al. 2016; Fuller et al. 2006; Latasa et al. 2010). Significant contributions by cryptophytes has also been observed in open ocean waters of the NW Mediterranean at the termination of the spring bloom (Vidussi et al. 2000) where they even dominated the surface mixed layer community at highly stratified stations. Interestingly, the higher contribution of cryptophytes in our study occurred towards the end of the spring bloom (TAN1212)(Figure S16), coincident with strong surface stratification and biomass accumulation (Chiswell et al. 2019), and supporting the importance that stratification may have on this group compared to salinity. *G. cryophile* and other species of *Teleaulax* have been reported as mixotrophic (Schneider et al. 2020), which could favor their increase in later stages of the spring bloom when the coincident decrease of nutrients and increase of potential preys tend to favor mixotrophy (Mitra et al. 2014).

#### 4.3.9. Heterotrophic and mixotrophic protists below the euphotic zone

We used the samples collected during six voyages of the Biophysical Moorings time-series between 2009-2012 to investigate the protistan community composition below the euphotic zone (Table 1). Metabarcode datasets here were clearly dominated by Radiolaria (Figure 7), a holoplanktonic amoeboid group with widespread distribution in modern oceans (Biard et al. 2016; Suzuki and Not 2015). Radiolaria are mainly heterotrophic protists with many mixotrophic species in the photic zone bearing endosymbiotic microalgae that can contribute substantially to primary production in oligotrophic oceans (Caron et al. 1995; Decelle et al. 2015). While Radiolaria are found throughout the entire water column, their contribution to plankton biomass (Biard et al. 2016; Boltovskoy and Correa 2016) and metabarcodes (Ollison et al. 2021) tends to be greater in the mesopelagic ocean, in a way consistent with our metabarcoding results. In this study, significant contributions of Radiolaria were mainly constrained to the aphotic zone (Figure 7). This depth-related pattern, contrasts with previously reported abundance of Radiolaria, and symbiotic Collodaria, in the sunlit ocean (Vargas et al. 2015). We found substantial contributions of photosymbiotic Collodaria at times, particularly in SAW (Figure S9), but the vertical distribution of Radiolaria below the euphotic zone suggested they were mainly composed of heterotrophic species. The high copy number of 18S rDNA in Radiolaria may contribute to their high relative abundance metabarcode datasets ((Gutierrez-Rodriguez et al. 2019; Vargas et al. 2015)); however, it is unlikely to be responsible for their dominance, particularly in relation to dinoflagellates and ciliates, which are also known to have high copy numbers (Gong et al. 2013; Piredda et al. 2017). Moreover, the positive relationship between cell length and 18S rDNA copy number (Biard et al. 2017; Zhu et al. 2005) and the higher C and N density (mass: volume) of Radiolaria compared to other protist (Mansour et al. 2021) suggest that gene-based relative abundance of these groups was likely reflected in their relative contribution to the community biomass.

Utilising their sticky pseudopodia and large size, Radiolaria dwelling below the euphotic zone can effectively intercept sinking particles and act as flux-feeders influencing the quality and quantity of vertical fluxes (Ohman et al. 2012; Stukel et al. 2019). The presence of mineral skeletons, made of silica (Polycystinean groups) or strontium sulfate (e.g. Acantharea), provides substantial mineral ballast (Takahashi 1983) conferring them a key role in vertical export that is supported by their common presence in sediment traps (Bernstein et al. 1987; Gutierrez-Rodriguez et al. 2019; Michaels et al. 1995; Preston et al. 2020) and their enrichment in suspended and sinking particles (Duret et al. 2020). Despite their abundance and their important role in biogeochemical processes (Biard et al. 2016; Guidi et al. 2016) little is known about the vertical distribution of Radiolaria particularly in the meso- and bathypelagic ocean (>500 m) (Biard and Ohman n.d.; Boltovskoy 2017; Llopis Monferrer et al. 2021; Ollison et al. 2021). In our study, we found an opposite distribution between Acantharea and RAD-B, which showed preference for the upper (<500 m) and deeper (>500 m) mesopelagic samples, respectively (Figure 7). Among Acantharea, most sequences were assigned to the order *Chaunchantida* (Figure S9, Figure S14), which has been found in sinking particles collected in the Southern Ocean (Duret et al. 2020) and the California Current (Gutierrez-Rodriguez et al. 2019; Preston et al. 2020). In a recent study conducted in the Southern Ocean, RAD B was reported to be enriched in small (<10 *µ*m) suspended particles relative to sinking particles (Duret et al. 2020) while RAD-A were consistently found in sinking particles reaching abyssal depths in the California Current Ecosystem (Preston et al. 2020). The higher contribution of RAD-B relative to RAD-A we found in our study is consistent with the tendency of RAD-B to remain suspended compared to RAD-A with greater sinking potential. Among the polycystinean Radiolaria, Spumellarida was the most important order, in a way consistent with observed abundance of this order in sinking particulate organic matter collected in sediment traps deployed in mesopelagic and abyssal depths in the California Current Ecosystem (Gutierrez-Rodriguez et al. 2019; Preston et al. 2020). Several ASVs assigned to Spumellarida Group I were among the most abundant species in our study (Figure S13). Interestingly, some of these ASVs showed preference for SAW while others were more abundant in STW, highlighting the need to improve our taxonomic knowledge of this group.

Relative contributions of ciliates were also higher below the euphotic zone, mainly driven by ASVs affiliated with Spirotrichea class (Grattepanche et al. 2016). Most abundant ASVs assigned to the order Strombidiida (Oligotrichia, Spirotrichea) and *Leegaardiella* sp. (Choreotrichia, Spirotrichea), which have been reported below the euphotic zone at meso- and bathy-pelagic depths (Duret et al. 2020; Grattepanche et al. 2016). In addition to Spirotrichea, class Oligohymenophorea and Nassophorea contributed substantially in both STW and SAW, mainly sustained by ASVs assigned to OLIGO5 (*Oligophymenophorea*) and *Discotrichidae* (*Nassophorea*), both previously reported in mesopelagic waters (Duret et al. 2020). The dominance of these groups was consistent across different water masses off eastern New Zealand although some species like *Leegaardiella* (Oligotrichia) and *Strombidium_k*_sp (Choreotrichia) showed preference for STW and SAW, respectively (Figure S11). Ciliates below the euphotic zone feed on bacteria and small protists associated with particulate organic matter (Caron et al. 2012). Several species also have the potential to engage in photoautotrophy and phagotrophy (Leles et al. 2017). By doing so, ciliates play an important role within planktonic food webs contributing to trophic transfer and nutrient recycling in the dark ocean. The high taxonomic diversity and abundance of heterotrophic protists in this and previous studies (Duret et al. 2020; Grattepanche et al. 2016; Ollison et al. 2021; Zoccarato et al. 2016) highlights their importance in planktonic systems below the euphotic zone and emphasises how little we know about their ecological role in the food web functioning. Further studies focusing on the taxonomic and functional diversity of heterotrophic protists have the potential to shed light on the trophic and biogeochemical processes that transform organic matter in the dark ocean, and hence significantly improve our understanding of the biological carbon pump and natural deep-ocean carbon sequestration.

## 5. CONCLUSIONS

The spatial diversity patterns observed are in agreement with global trends of decreasing diversity at higher latitudes and colder waters. However, deviations from this general pattern were also observed regionally. Species richness and diversity of protist communities in the STF were systematically lower compared to adjacent STW and SAW waters in the northern and southern regions of the STFZ surveyed, highlighting the importance of oceanographic features in determining regional diversity. Dinoflagellates and green algae co-dominated the protist community in the euphotic zone but water-mass specificity emerged at lower taxonomic levels within these and other major taxonomic groups and the community composition varied consistently between water masses. Within green algae for instance, *Mamiellophyceae* class dominated in STW driven by several species showing different regional abundance, while *Chloropicophyceae* class became dominant in SAW where several ASVs assigned to *Chloroparvula pacifica* appeared among the most abundant taxa. Interestingly, other less abundant ASVs identified as *Chloroparvula pacifica* showed statistically significant preference for STW. Analogous intra-specific variability was observed within species belonging to other phytoplankton classes with widespread distribution (e.g. *P. antarctica*, prymnesiophytes; *P. calceolata*, pelagophytes) suggesting the genotypic diversity may be linked to ecological traits that influence distribution patterns. Although chl *a* levels comprised in this study were relatively low, small rather than large taxa dominated the phytoplankton proliferations associated with spring bloom conditions and the STFZ suggesting that picoplankton can also be important for primary and export production either directly or through zooplankton grazing. The mesopelagic zone was clearly dominated by radiolarian sequences, supporting the importance of this group for the functioning of the dark ocean. Taxonomic assignation revealed a diverse assemblage of Radiolaria and a taxon-specific water mass and vertical distribution patterns. However, further research is needed about the ecology of these organisms to link this compositional variability to their function in the system.

## 7. ACKNOWLEDGEMENTS

We acknowledge the crew of RV *Tangaroa* and RV *Kaharoa* for their efforts in facilitating the sampling throughout all the voyages included in this work. We thank Els Maas and Cliff Law for collecting DNA samples during KAH1303 voyage and for sharing the physical and chemical data obtained in. this voyage. Thank you to Daniel Vaulot for critical feedback on PR2 references and assignation. We are thankful to the anonymous reviewers for their efforts and constructive comments that helped improve the manuscript. We are grateful to the Royal Society of New Zealand which funded this research through the Catalyst Seeding General fund (grant reference number 16-NIW-009-CSG) and foster the collaboration between the Station Biologique de Roscoff and NIWA. This work is also supported by NIWA via the New Zealand Ministry of Business, Innovation and Employment’s Strategic Science Investment Funding to the National Coasts Oceans Center. ALS was supported by the Singapore Ministry of Education, Academic Research Fund Tier 1 (RG26/19).

## 8. AUTHOR CONTRIBUTION STATEMENT

- Andres Gutiérrez-Rodríguez: Conceptualization, Methodology, Formal analysis, Investigation, Data curation, Visualization, Writing - Original Draft & Funding acquisition
- Adriana Lopes dos Santos: Conceptualization, Methodology, Investigation, Data curation, Writing - Original Draft & Funding acquisition.
- Karl Safi: Investigation, Writing - Review & Editing.
- Ian Probert: Conceptualization, Writing - Review & Editing and Funding acquisition.
- Fabrice Not: Conceptualization, Writing - Review & Editing and Funding acquisition.
- Denise Fernandez: Conceptualization, Writing - Review & Editing
- Jaret Bilewitch: Methodology, Writing - Review & Editing
- Debbie Hulston: Methodology, Writing - Review & Editing
- Matt Pinkerton: Writing - Review, Editing & Funding acquisition.
- Scott D Nodder: Conceptualization, Investigation, Data curation, Writing - Original Draft & Funding acquisition.

## 9. SUPPLEMENTARY MATERIAL

**Table. S1.** Supplementary Table S1. Table with the number of samples, the median number of reads per sample and the number of ASVs in the entire dataset and within each water mass.

| Water mass | N samples | N reads (median) | N ASVs |
| --- | --- | --- | --- |
| **All** | **482** | **15913** | **16861** |
| SAW | 120 | 17158 | 7377 |
| STF | 91 | 16431 | 4261 |
| STW | 271 | 15630 | 12407 |

**Table. S2.** Supplementary Table S2. Summary of PERMANOVA analysis results conducted to calculate the significance of environmental variables ability to explain ASV composition on a subset of 188 samples collected from STW (n=34), STF (n=53), SAW (n=95) for which measurements of presented variables were available. Analysis was conducted with the Adonis function of the vegan R package.

| Variable | Df | SumsOfSqs | MeanSqs | F.Model | R2 | Pr(>F) |
| --- | --- | --- | --- | --- | --- | --- |
| Temperature | 1 | 4.929 | 4.9288 | 21.639 | 0.08489 | 0.001 |
| Salinity | 1 | 7.831 | 7.8308 | 34.38 | 0.13487 | 0.001 |
| median NO<sub>3</sub> | 1 | 3.362 | 3.3617 | 14.759 | 0.0579 | 0.001 |
| median Chla | 1 | 1.625 | 1.6246 | 7.133 | 0.02798 | 0.001 |
| Residuals | 177 | 40.316 | 0.2278 |  | 0.69436 |  |
| Total | 181 | 58.062 |  |  | 1 |  |

**Fig. S1.**
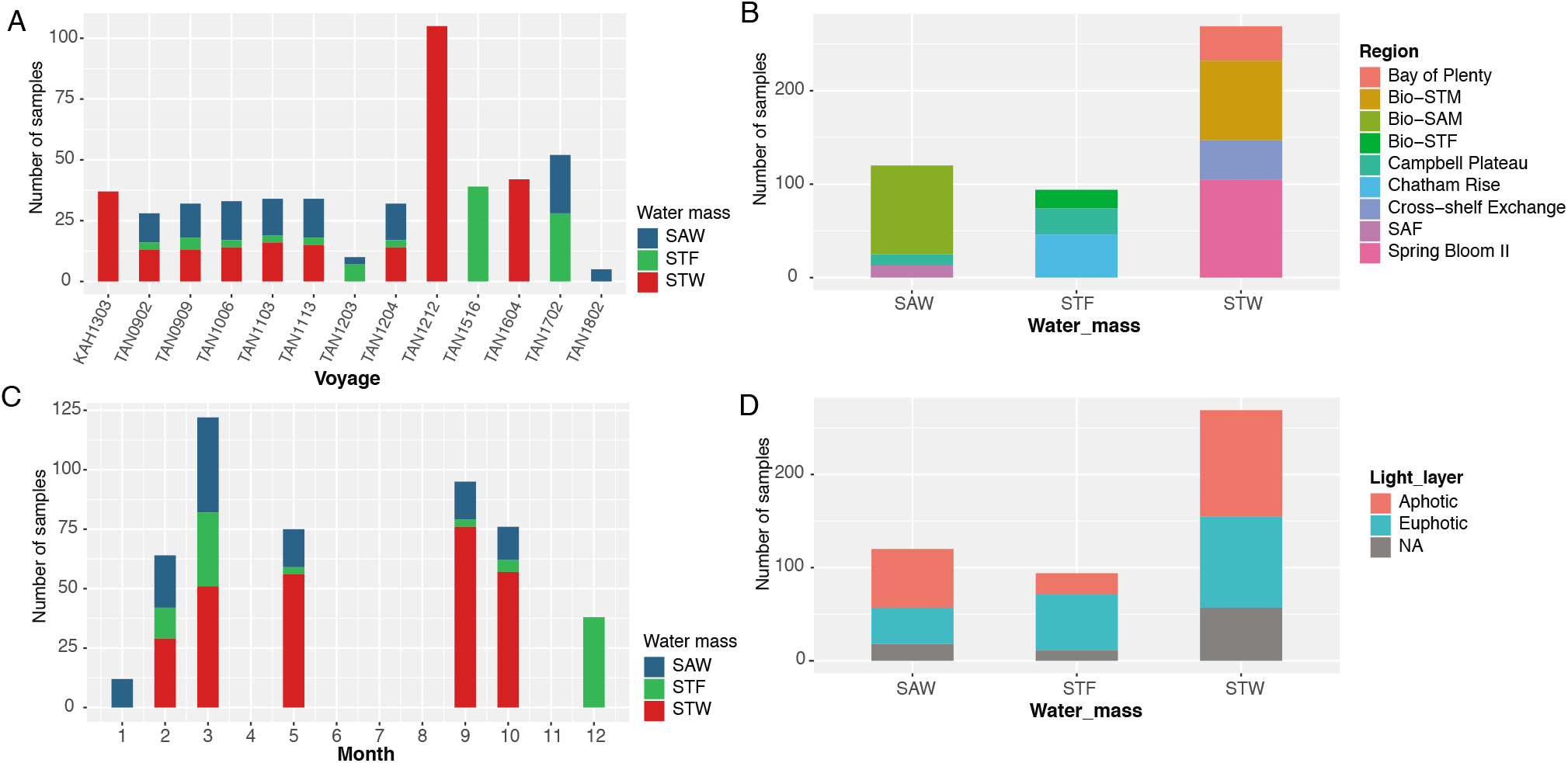
Supplementary Figure S1. Number of DNA samples from each water mass surveyed in different (A) cruises, (B) regions, (C) months of the year, and (D) photic zone.

**Fig. S2.**
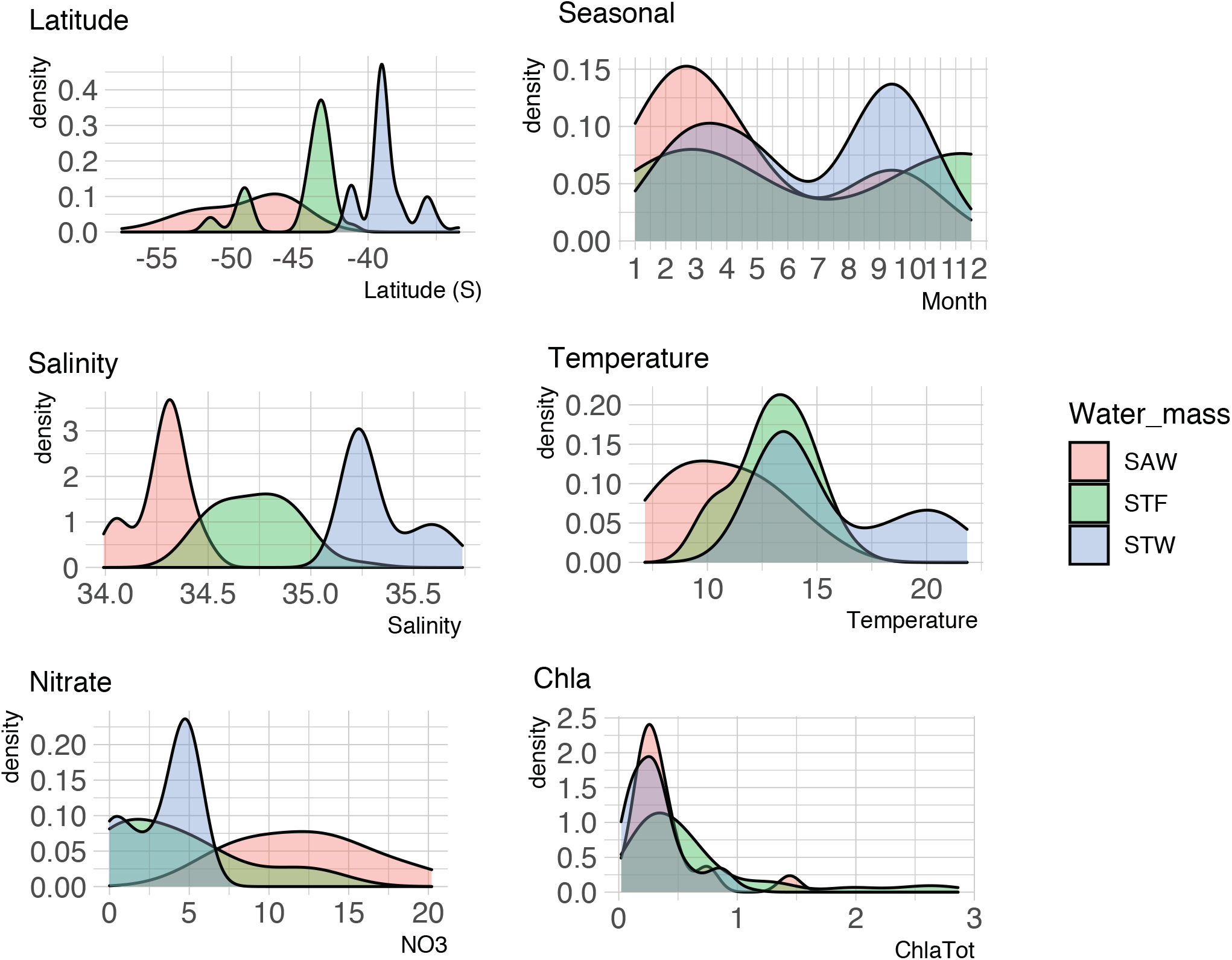
Supplementary Figure S2. Density distribution of DNA samples across latitude, season, and mixed-layer temperature, salinity nitrate, and chlorophyll *a* concentrations in each water mass surveyed.

**Fig. S3.**
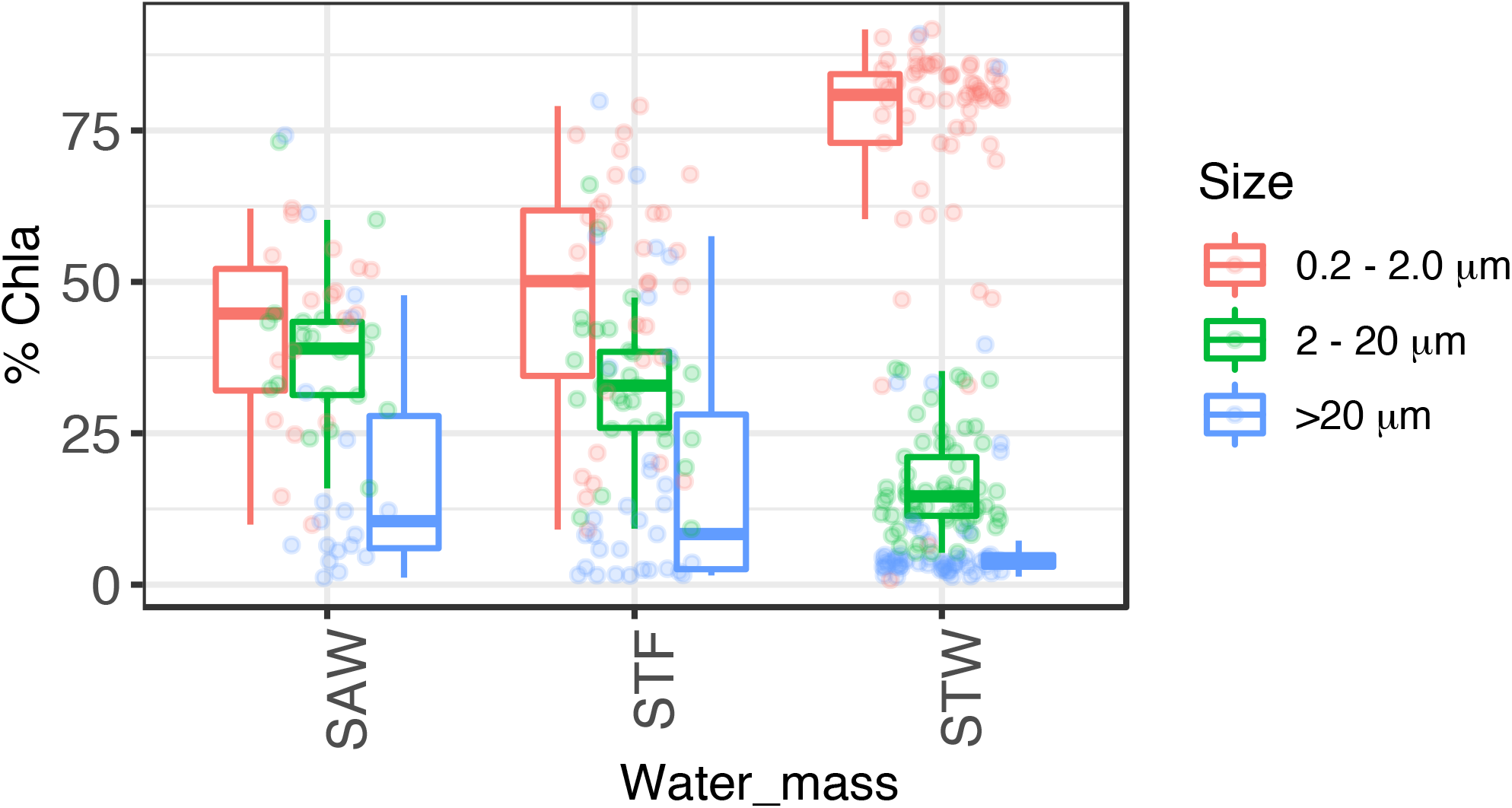
Supplementary Figure S3. Phytoplankton community size-structure. Box-plots show the chlorophyll a concentration associated to pico-, nano-, and micro-phytoplankton nominal size fractions (0.2-2 *µ*m, 2-20 *µ*m, >20 *µ*m) in the surface mixed-layer of each water masses. Each dot correspond to a single sample. Box-plots show the median, first and third quartiles and the values within the *±*1.5 *∗ IQR* (IQR, interquartile range). The figure includes data from TAN1516/Chatham Rise, TAN1702/Campbell Plateau, TAN1212/Spring Bloom, TAN1203/SOAP; n = 102.

**Fig. S4.**
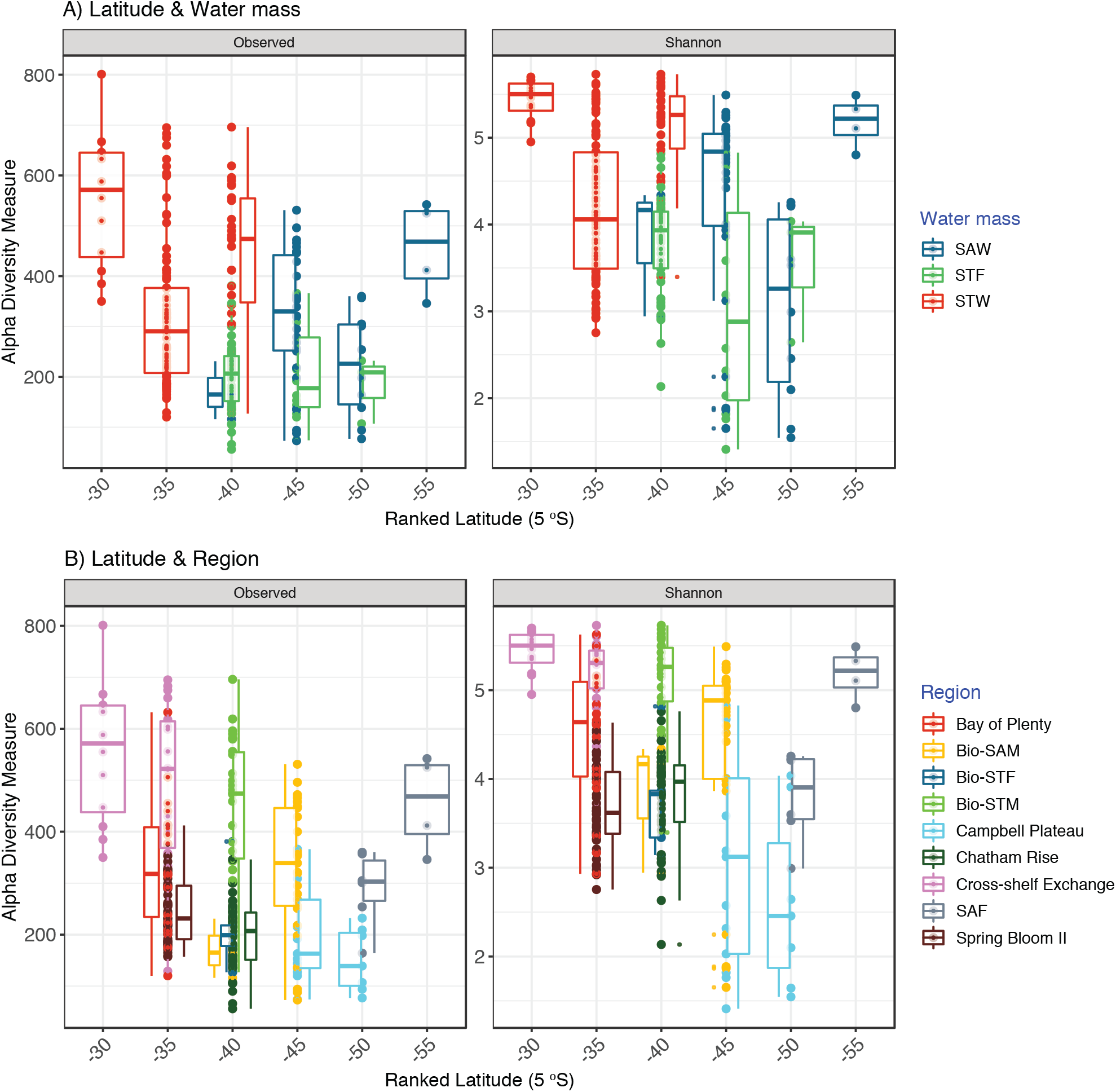
Supplementary Figure S4. Protist diversity richness and Shannon diversity index binned in 5° latitude bins and color coded by (A) water mass and (B) regions.

**Fig. S5.**
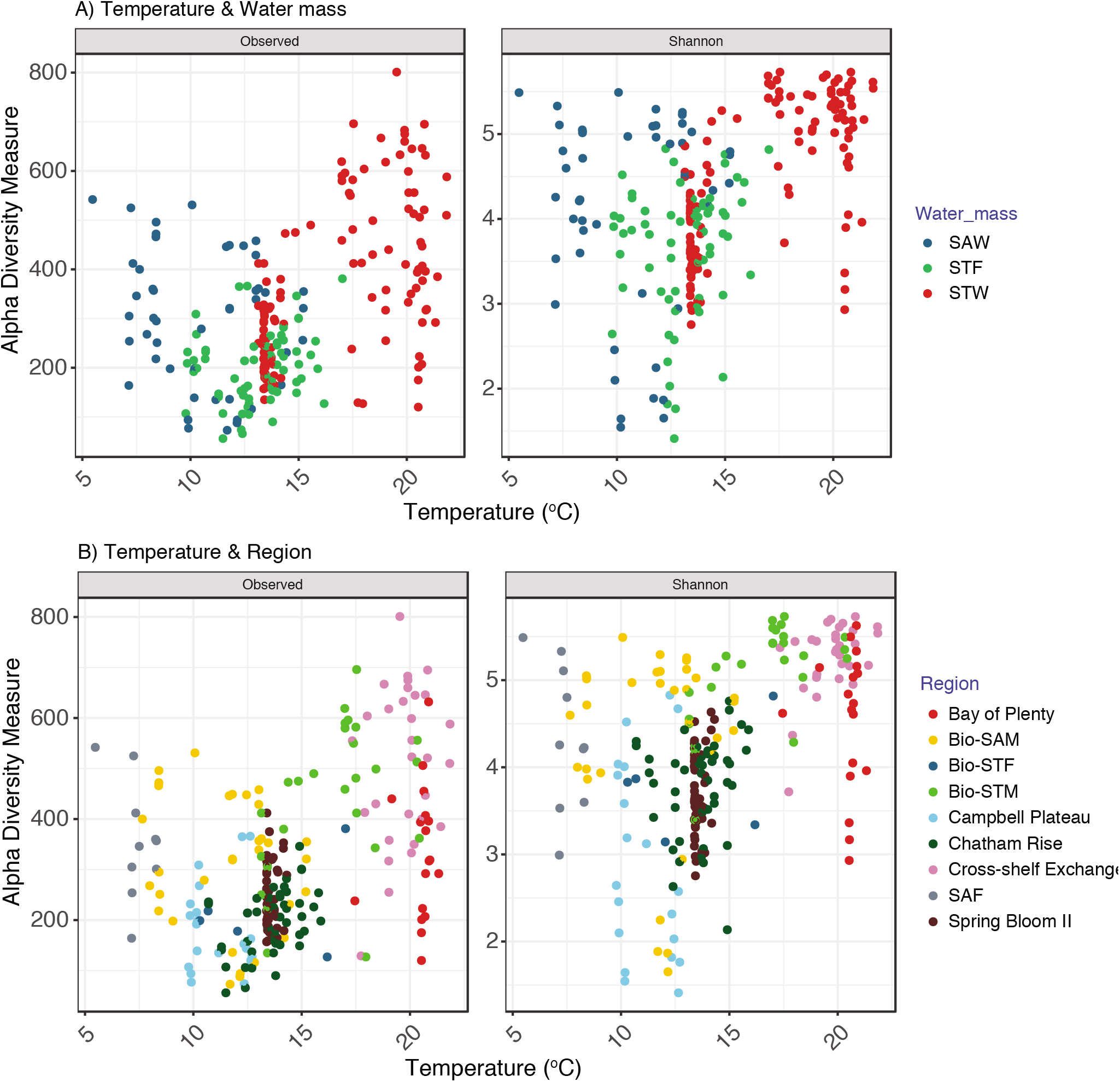
Supplementary Figure S5. Diversity and temperature. Protist diversity richness and Shannon diversity index binned in 5° latitude bins and color coded by (A) water mass and (B) regions.

**Fig. S6.**
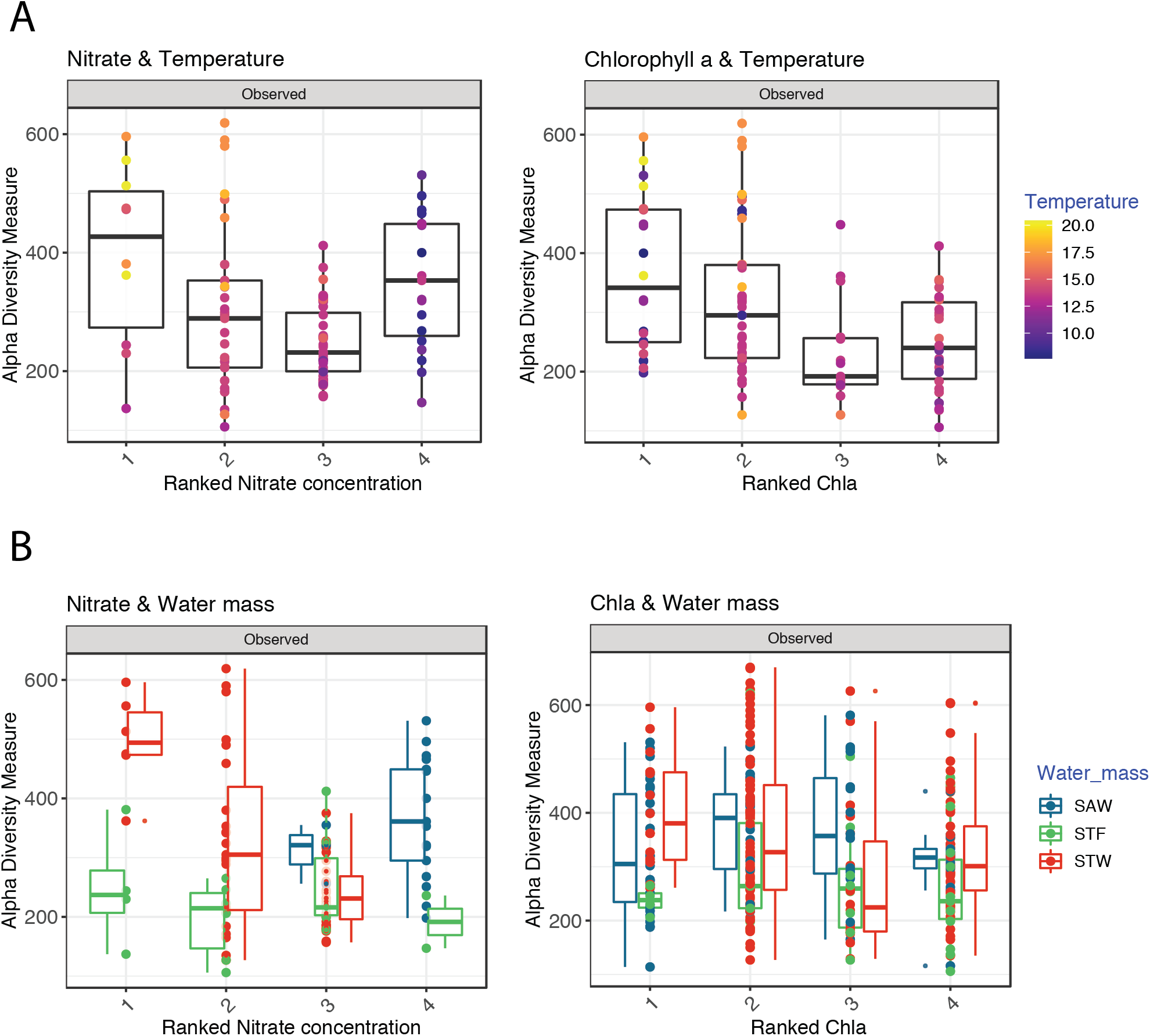
Supplementary Figure S6. Diversity and trophic conditions. Mean protist diversity richness in relation to nitrate and chlorophyll a concentrations in the euphotic zone and binned in four ranks and color coded by (A) temperature and (B) water mass. Function ntile was used to break individual measurements into buckets of the same size.

**Fig. S7.**
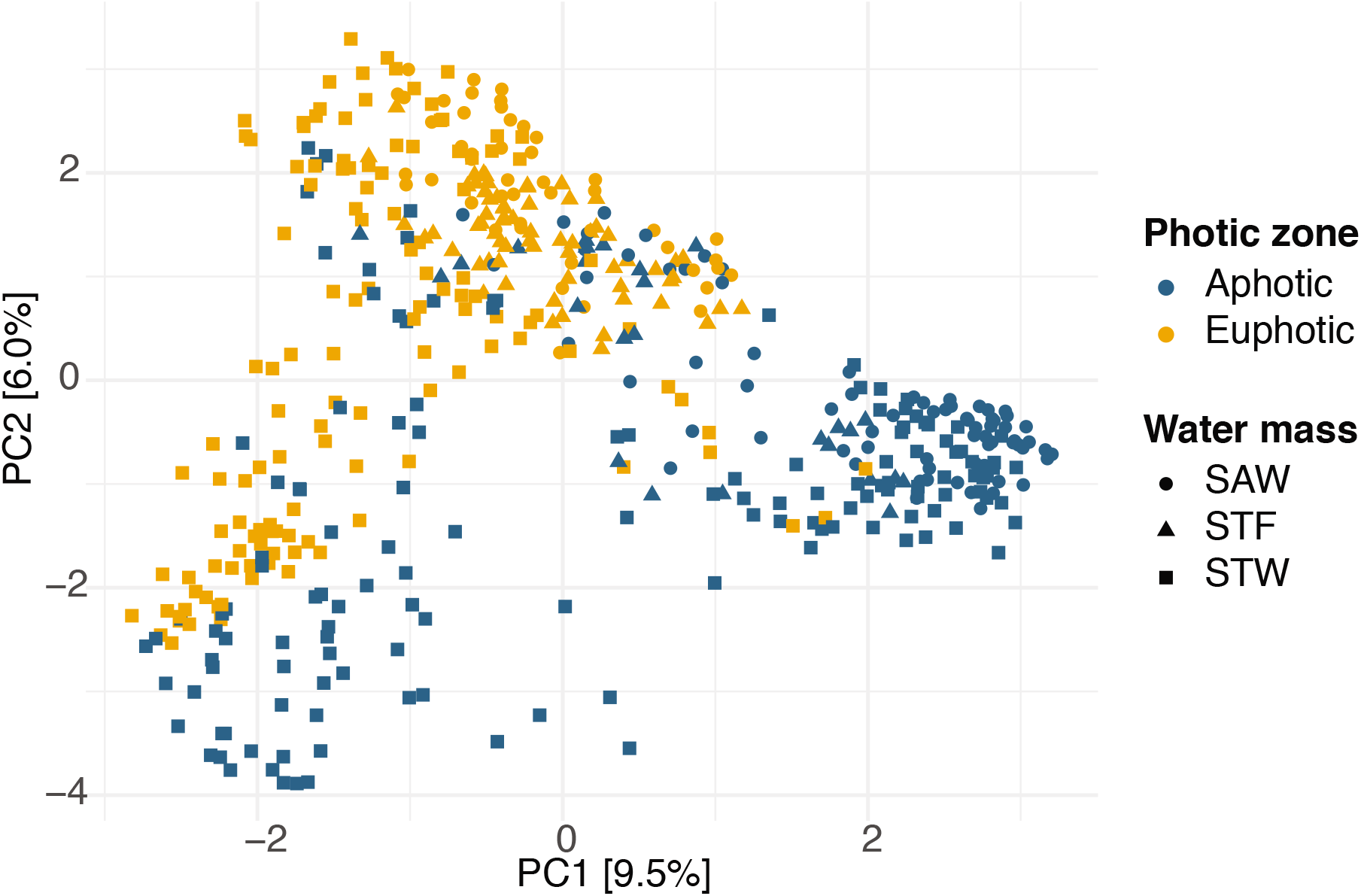
Supplementary Figure S7. Principal component analysis based on ASV composition of all samples coded by symbol color and shape, respectively, for the light layer and water masses where the samples were collected from.

**Fig. S8.**
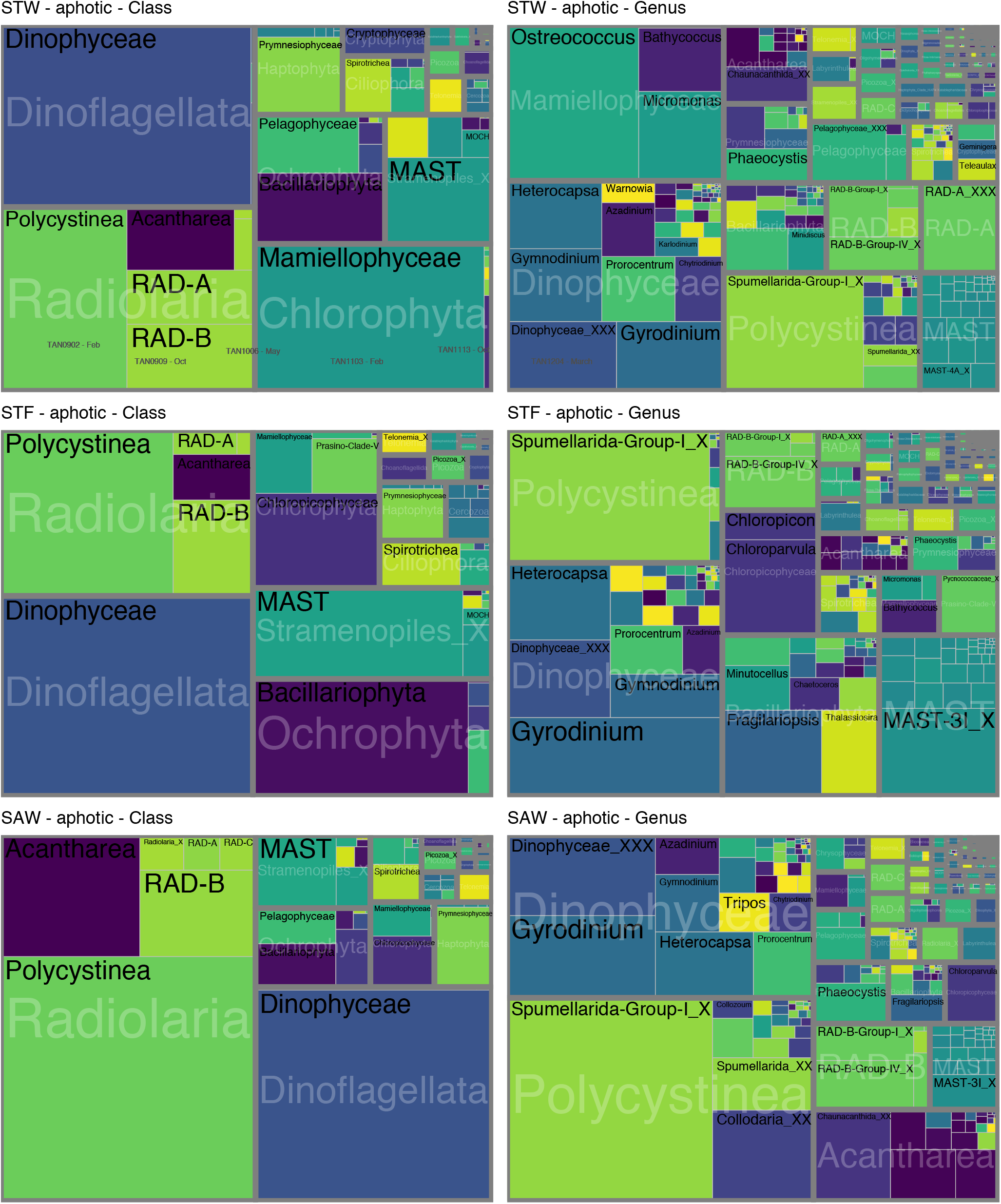
Supplementary Figure S8. Treemaps showing the mean relative abundance of main protistan groups divisions, class and genus in the aphotic zone of the STW, STF, and SAW sites. The area of each taxonomic group in the treemap represents the read abundances affiliated to each group standardized to the median sequencing depth across samples [median sum otus * (otu reads / sum (otu reads)]

**Fig. S9.**
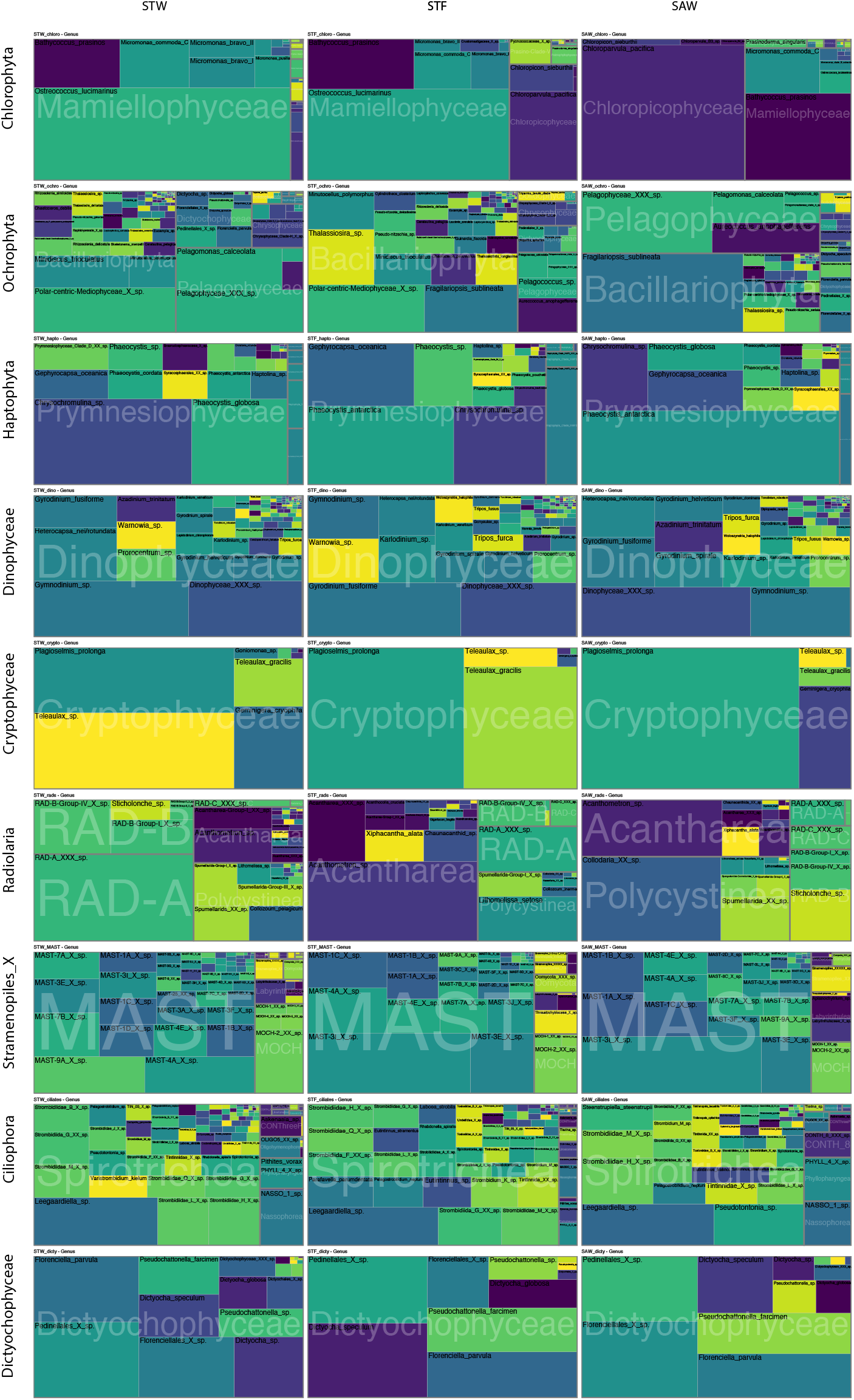
Supplementary Figure S9. Treemaps showing the mean relative abundance of main protistan classes and species within major taxonomic divisions in the euphotic zone of STW, STF, and SAW water masses.

**Fig. S10.**
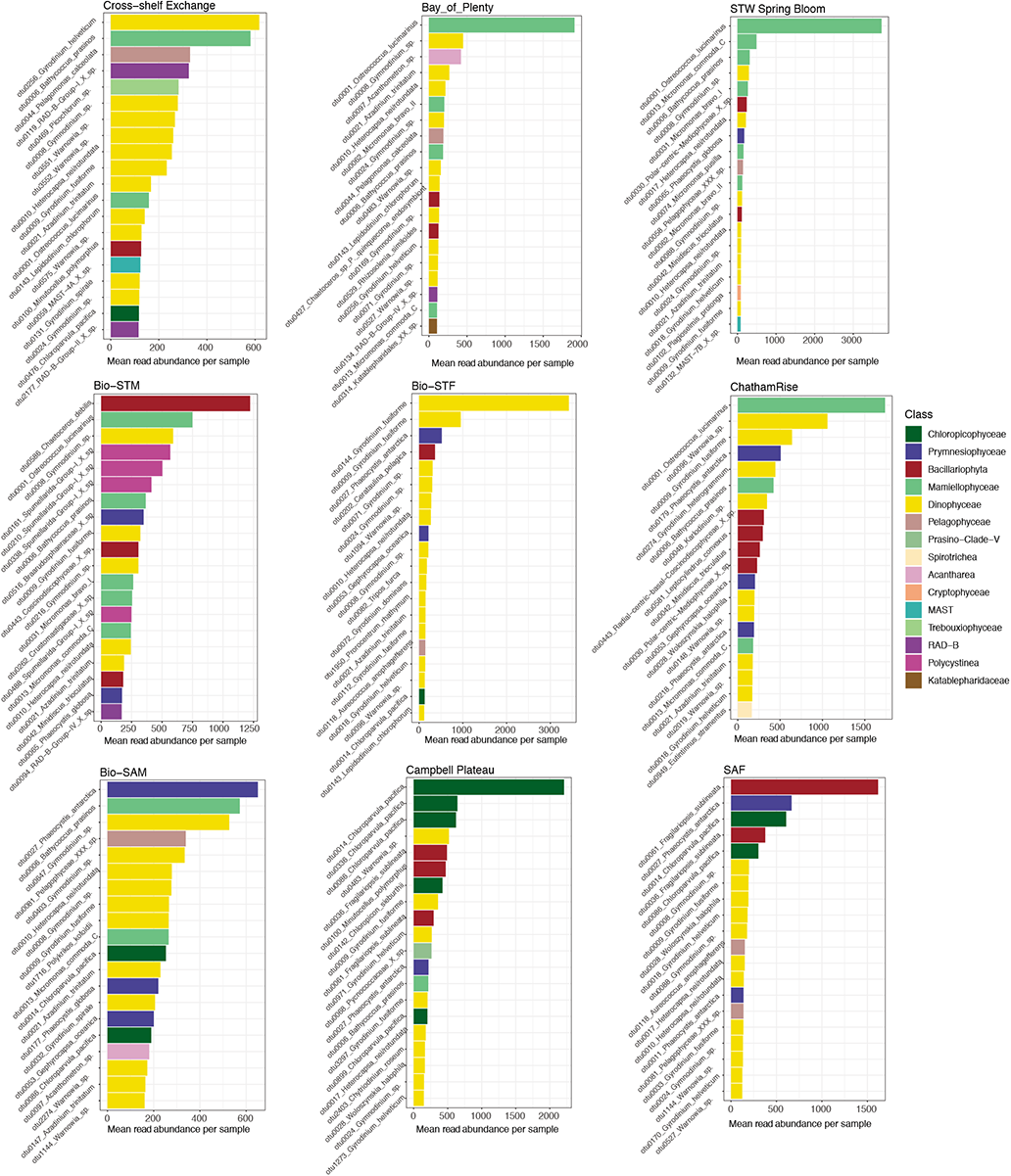
Supplementary Figure S10. Mean standardized read abundance of most abundant ASVs and assigned species in the euphotic zone of different regions and voyages, color coded for their class affiliation.

**Fig. S11.**
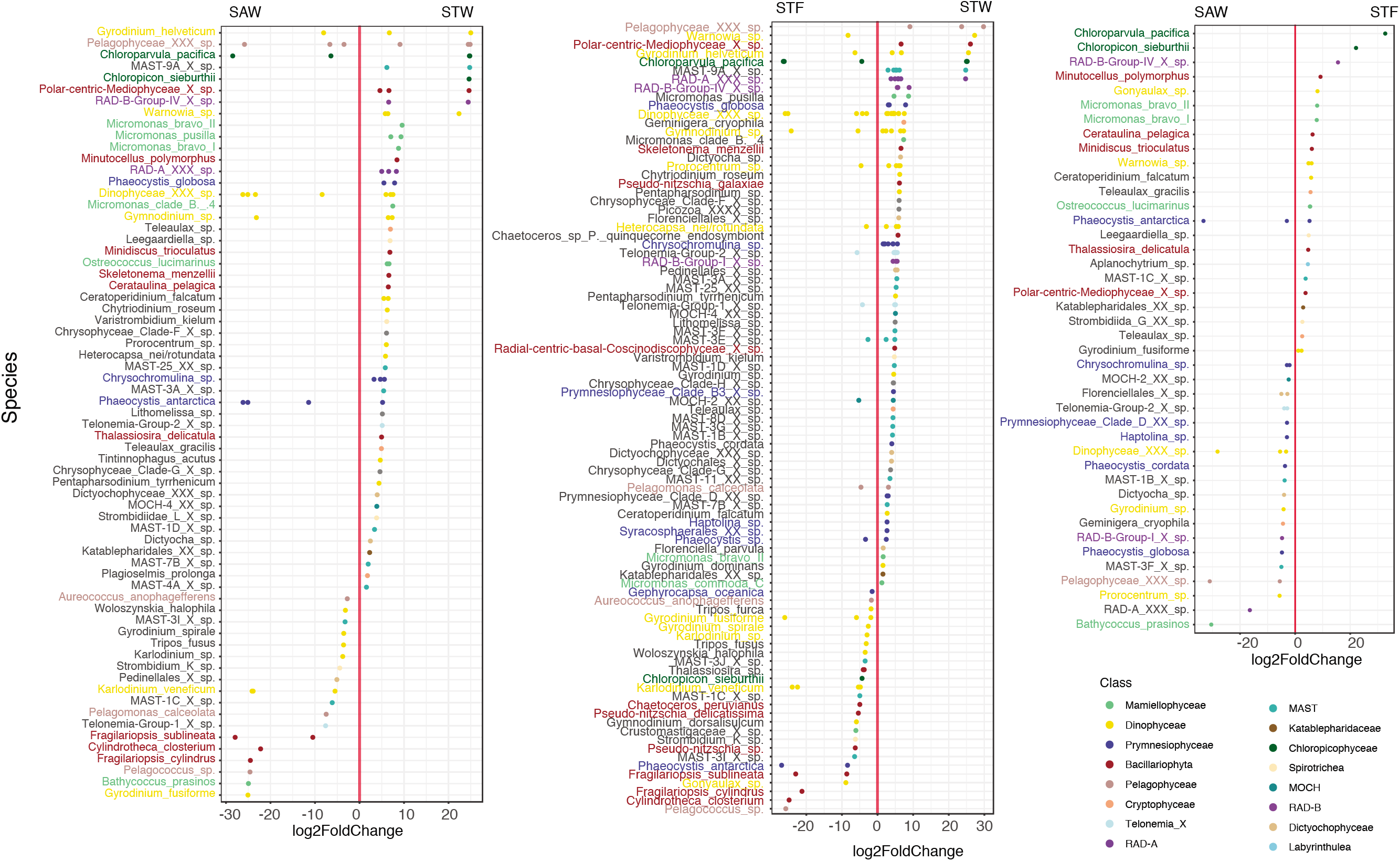
Supplementary Figure S11. Results from DESeq2 analysis depicting the species (Y-axis) with significantly different distribution between the euphotic zone of STW, SAW and STF waters. Difference in the distribution is expressed as the log2 fold change of the difference (X-axis). Each dot correspond to a different ASV color coded by their class affiliation.

**Fig. S12.**
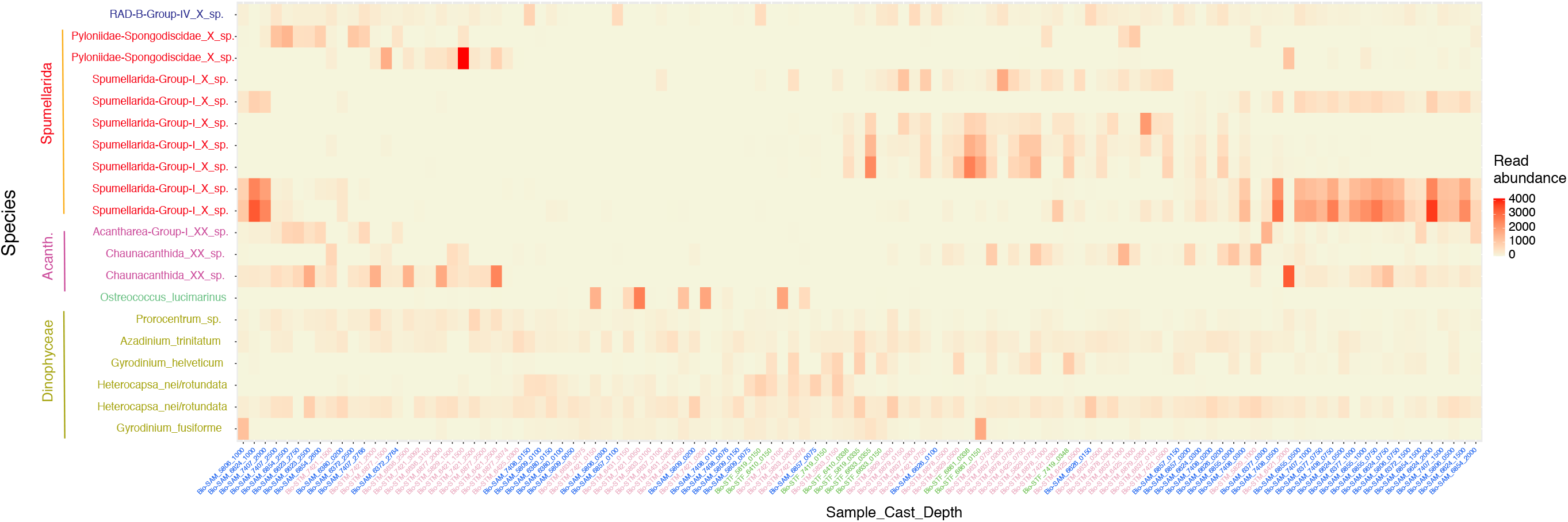
Supplementary Figure S12. Heatmap showing the abundance patterns of the top20 most abundant species in the aphotic zone of the Biophysical Mooring samples. Read abundance were normalized to mean sequencing depth. Samples are clustered using nMDS and Jaccard distance, and color coded according to the sampling site they were collected from (Bio-STM, purple; Bio-STF, green; Bio-SAM, blue). Species were organized and color coded by class affiliation.

**Fig. S13.**
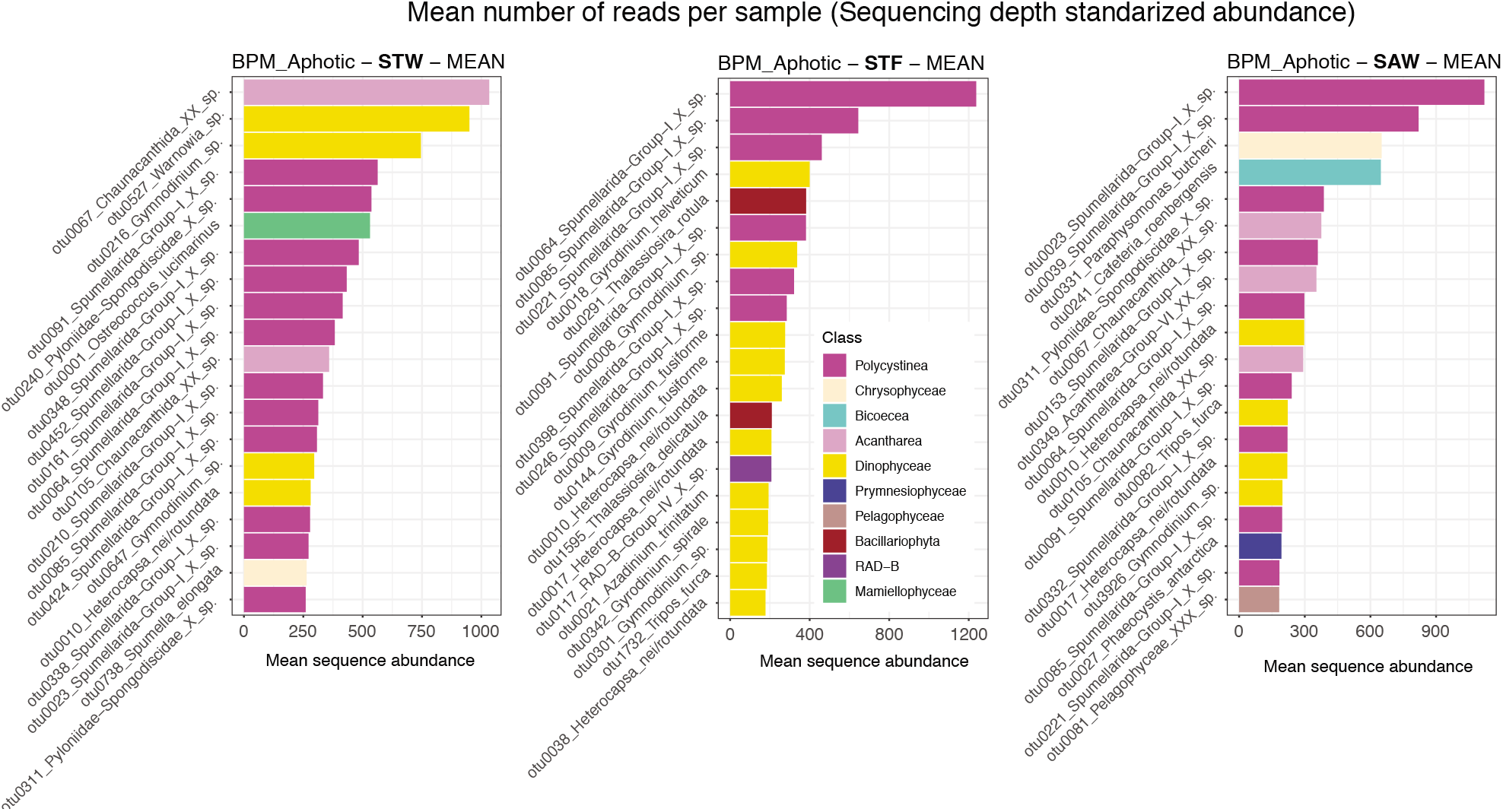
Supplementary Figure S13. Mean standardized read abundance of most abundant ASVs and assigned species in the aphotic zone of different water masses. Bars corresponding to each species color coded for their class affiliation.

**Fig. S14.**
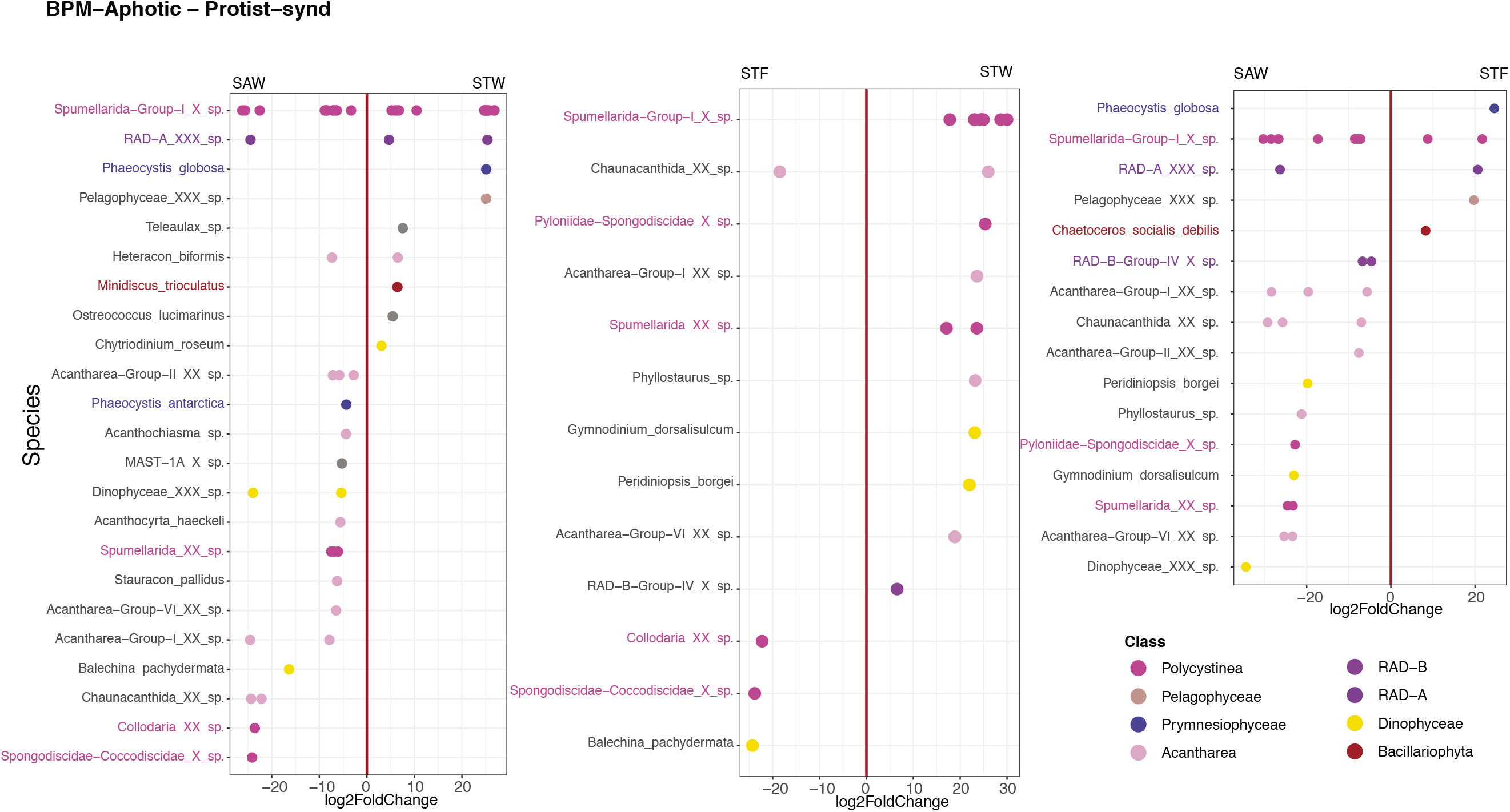
Supplementary Figure S14. Results from DESeq2 analysis depicting the species (Y-axis) with significantly different distribution between the aphotic zone of STW, SAW and STF waters. Difference in the distribution is expressed as the log2 fold change of the difference (X-axis). Each dot correspond to a different ASV color coded by their class affiliation.

**Fig. S15.**
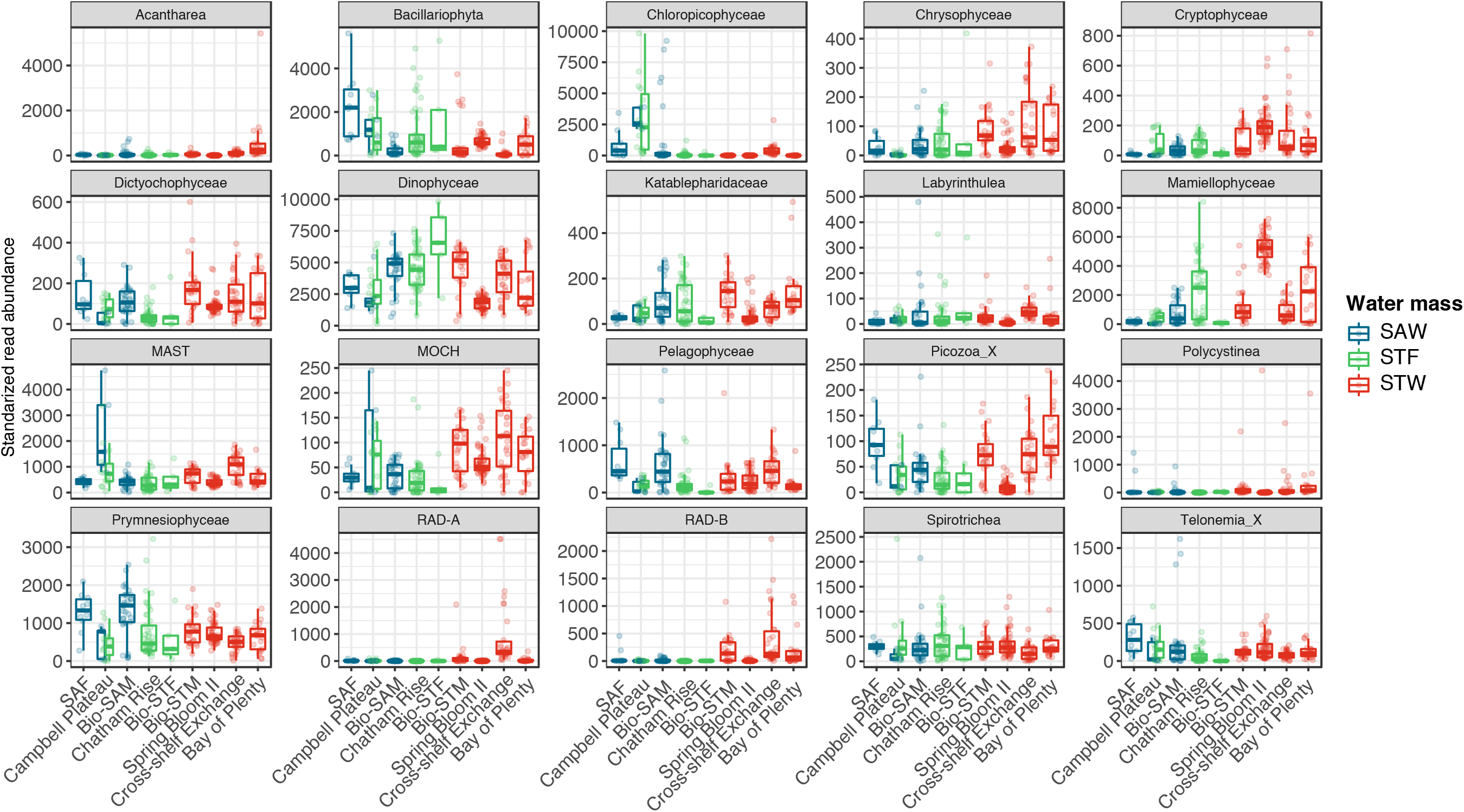
Supplementary Figure S15. Box-plots showing standardized read abundance of twentieth most abundant protist classes in the euphotic zone of different regions and color coded by different water masses. Box-plots show the median, the first and third quartiles (lower and upper hinges) and the values within the *±*1.5 *∗ IQR* (IQR, interquartile range) (line). Points represent values of single samples

**Fig. S16.**
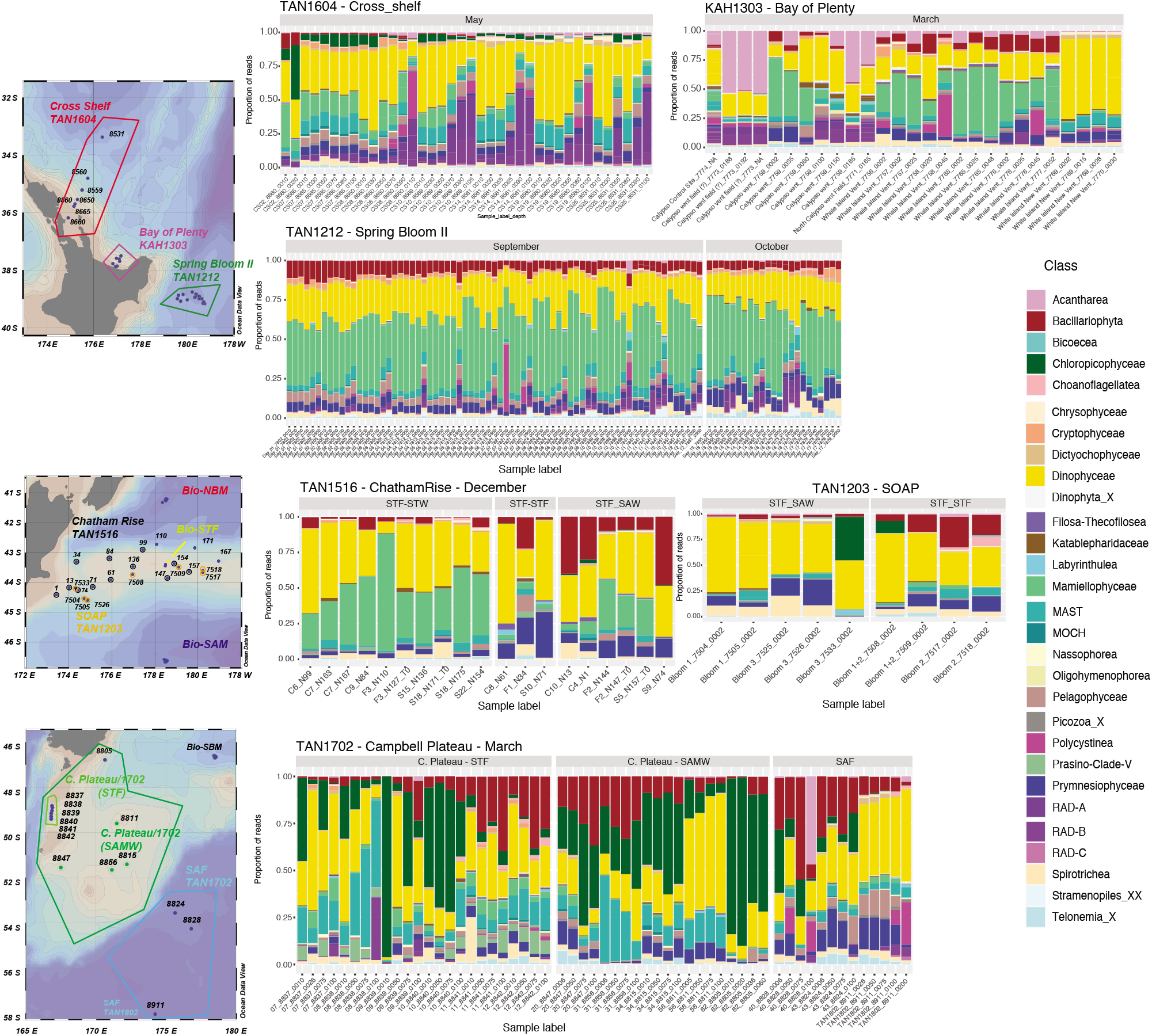
Supplementary Figure S16. Relative read abundance of main protistan classes in samples collected during different voyages and regions within the STW, STF and SAW water masses.

**Fig. S17.**
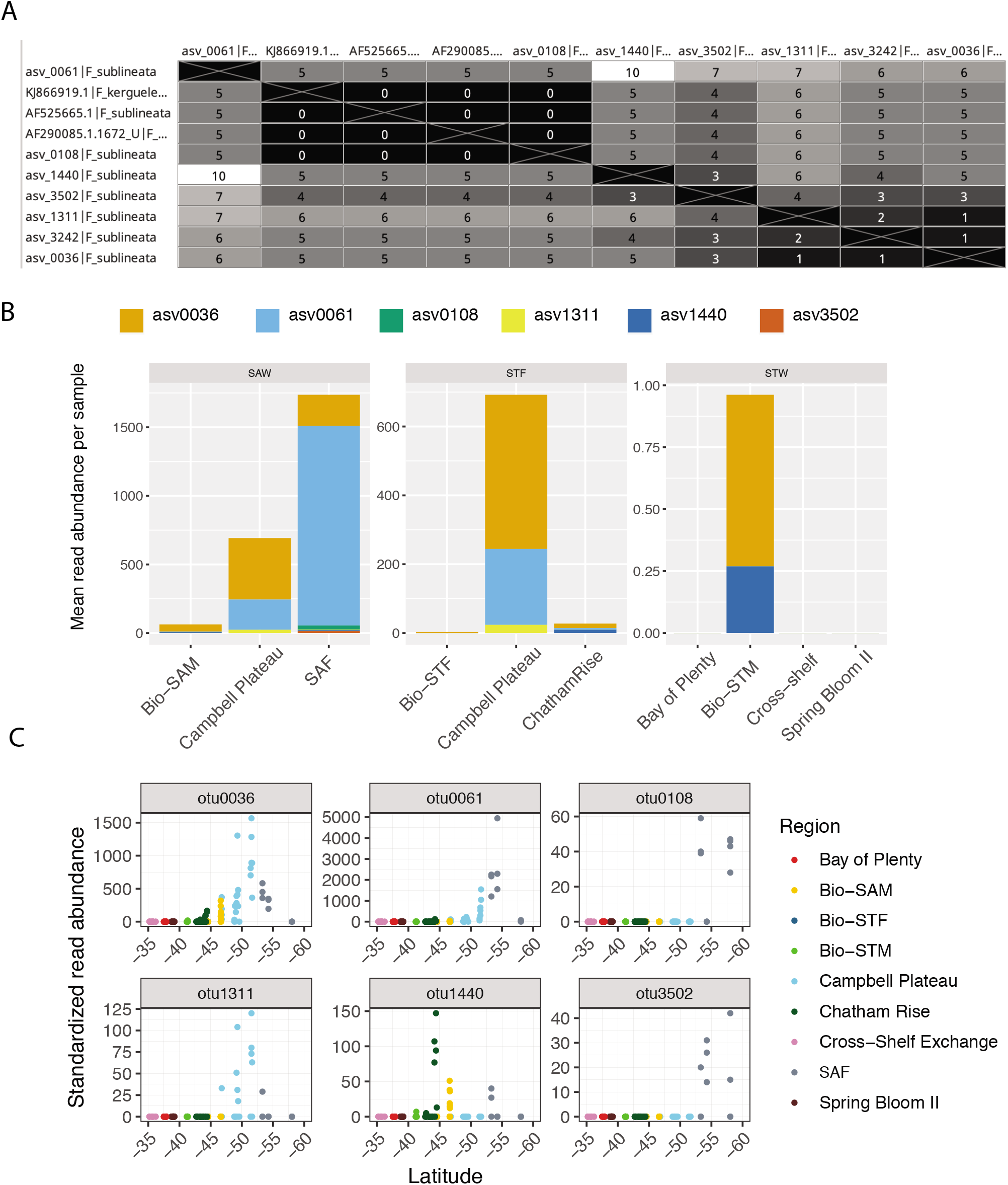
Supplementary Figure S17. (A) Matrix showing the number of nucleotide mismatches in the v4 region of the 18S rRNA between the most abundant ASVs assigned to *F. sublineata* and the annotated sequences for this and *F. kerguelensis* species included in PR2 reference database. (B) mean relative abundance of these ASVs in the euphotic zone of different water masses and regions. (C) distribution of these ASVs abundance (standardized read abundance) in each euphotic sample in relation to latitude and color coded by region.

